# Regulation of Auditory Sensory Neuron Diversity by Runx1

**DOI:** 10.1101/2022.08.02.502556

**Authors:** Brikha R Shrestha, Lorna Wu, Lisa V Goodrich

## Abstract

Functional heterogeneity among sensory neurons is a cardinal property of the vertebrate auditory system, yet it is not known how this heterogeneity is established to ensure proper encoding of sound. Here, we show that the transcription factor Runx1 controls the composition of molecularly and physiologically diverse sensory neurons (Ia, Ib, Ic) in the murine cochlea, which collectively encode a wide range of sound intensities. *Runx1* is enriched in Ib and Ic spiral ganglion neuron (SGN) precursors by late embryogenesis. Loss of *Runx1* from embryonic SGNs (*Runx1*^CKO^) shifted the balance of subtype identities without affecting neuron number, with more SGNs taking on Ia identities at the expense of Ib/Ic identities, as shown by single cell RNA-sequencing. This conversion was more complete for genes linked to neuronal function than for those related to connectivity. Accordingly, although synaptic position did not change, synapses in the Ib/Ic location took on Ia-like properties. Suprathreshold responses to sound were enhanced in the auditory nerve of *Runx1*^*CKO*^ mice, confirming an expansion of neurons behaving functionally like Ia SGNs. Fate-mapping experiments further showed that deletion of *Runx1* shortly after birth also redirected Ib and Ic SGNs towards Ia identity, indicating that SGN subtype identities remain plastic postnatally. Altogether, these findings show that diverse neuronal identities essential for normal auditory stimulus coding arise in a hierarchical fashion that remains malleable during postnatal development.

## Introduction

Across sensory systems, the burden of encoding stimuli is split across populations of related yet distinct sensory neuron subpopulations. These neurons differ at a functional level, for example, in terms of response sensitivity and input-output relationship, and such diversity constitutes a key circuit motif upon which a broad range of stimulus features are functionally mapped. For example, in the auditory periphery, Type I spiral ganglion neurons (SGNs) with different response thresholds collectively encode the wide range of sound intensities an animal may encounter (Bharadwaj et al., 2014; Rutherford et al., 2021; Shrestha and Goodrich, 2019). These neurons are classified based on transcriptomic profiles into three major subtypes (Ia, Ib, Ic) (Petitpré et al., 2018; Shrestha et al., 2018; Sun et al., 2018). Based on anatomy, these subtypes correspond to prior physiology-based groups with high-, medium-, and low-spontaneous rate (SR) groups that exhibit low-to-high thresholds, respectively, for sound-driven response (Liberman, 1978). Degeneration of Ic SGNs or loss of their synapses after sound overexposure (Furman et al., 2013) and in old age (Schmiedt et al., 1996; Shrestha et al., 2018) is thought to cause perceptual deficits, given the proposed significance of this SGN subtype for hearing against background noise (Costalupes, 1985; Kujawa and Liberman, 2015; Young and Barta, 1986) and of complex sounds (Bharadwaj et al., 2014). It is therefore believed that the full range of SGN subtype diversity is essential for high-fidelity auditory perception. However, it remains unclear how a stereotyped distribution of SGNs with physiologically distinct properties is established.

A recurring theme in neural development is the use of transcription factors (TFs) that act sequentially to produce functionally and morphologically distinct types of neurons (Allan and Thor, 2015; Hobert, 2011; Shirasaki and Pfaff, 2002). Early acting TFs direct neuronal progenitors towards progressively more restricted fates, ultimately activating sets of TFs that induce cohorts of genes needed for mature function. For instance, the TF Gata3 steers inner ear neuronal progenitors towards an auditory fate and activates expression of effector TFs, such as Mafb, which controls the formation of synapses with inner hair cells (IHC) (Yu et al., 2013). In parallel, TFs likely act in a subtype-specific manner to generate Ia, Ib, and Ic SGNs that exhibit physiological and anatomical differences. For example, *Pou4f1* is expressed in a high-to-low gradient from Ic to Ib SGNs and is off in Ia neurons (Shrestha et al., 2018). Loss of *Pou4f1* in SGNs alters voltage dependence of Ca^2+^ influx as well as magnitude of Ca^2+^ entry in IHCs (Sherrill et al., 2019), suggesting a role in instructing presynaptic features, presumably via trans-synaptic signaling. However, other aspects of SGN subtype identity appear unchanged, leaving open the question of how SGN subtype heterogeneity is established. Ia, Ib, and Ic precursors are distinguishable based on their molecular profiles by late embryogenesis in mice (Petitpré et al., 2022). Subtype-specific patterns of gene expression continue to be refined during the first postnatal week (Shrestha et al., 2018), and individual neurons do not exhibit mature physiological properties and synaptic heterogeneity until the end of the first month (Liberman and Liberman, 2016). Further, the postnatal consolidation of SGN subtype identity is both flexible and activity-dependent, as more SGNs acquire Ia identities when IHC-SGN signaling is disrupted (Shrestha et al., 2018; Sun et al., 2018). Thus, additional TF(s) likely coordinate acquisition of subtype identity in the spiral ganglion, perhaps in response to activity or other signals in the environment.

An excellent candidate for SGN diversification is the transcription factor Runx1, which is detected at similar levels in Ib and Ic neurons but absent from Ia neurons (Shrestha et al., 2018). Likewise, Runx1 is restricted to subsets of developing dorsal root ganglion (DRG) neurons and promotes the non-peptidergic nociceptive identity within this population (Chen et al., 2006; Kramer et al., 2006). In this role, Runx1 acts both as transcriptional activator and repressor, regulating a battery of ion channel and receptor genes, i.e. genes that could impart functional differences among DRG neurons (Chen et al., 2006). Notably, in this system, Runx1 functions at the nexus of extrinsic (nerve growth factor-driven) and cell-intrinsic signals to regulate gene transcription associated with non-peptidergic identity (Huang et al., 2015). Furthermore, Runx1 regulates postmitotic cell fate decisions in several other cell types, with important roles in definitive hematopoiesis as well as specification of T lymphocytes in the thymus (Mevel et al., 2019).

Here, we investigated the role of *Runx1* in diversification of Type I SGNs in the mouse cochlea. We found that, upon ablation of *Runx1* function, Ib and Ic identities are significantly depleted without overt neuronal loss, resulting in overabundance of Ia identity. This change in the proportions of Ia, Ib, and Ic SGNs is consequential for sound encoding at the auditory nerve, as mutant animals exhibit heightened neural responses to suprathreshold stimuli. In addition, we show that *Runx1-*positive precursors are biased to become Ib/Ic SGNs by birth and that this expression must be maintained postnatally for Ib and Ic SGNs to hold on to their nascent identities. Together, these findings show that plasticity in neuronal identity contributes to neural circuit formation and identify *Runx1* as a key regulator of neural diversity in the auditory periphery subserving sound encoding.

## Results

To determine how Type I SGN diversity is generated, we considered transcription factors expressed differentially among Type I SGNs based on their single cell transcriptional profiles (Petitpré et al., 2018; Shrestha et al., 2018). *Runx1* stood out as a strong candidate given its enrichment in Ib/Ic SGNs (Fig. 1A, plotted based on data in (Shrestha et al., 2018)) and known roles in determining alternative cell fates among lymphocytes, skin cells, and DRG neurons. Since transcriptional identities of Type I SGNs begin to coalesce embryonically (Petitpré et al., 2018, 2022; Shrestha et al., 2018), we reasoned that if *Runx1* is involved in regulating SGN identities, it may exhibit segregated expression before birth.

**Figure 1:**
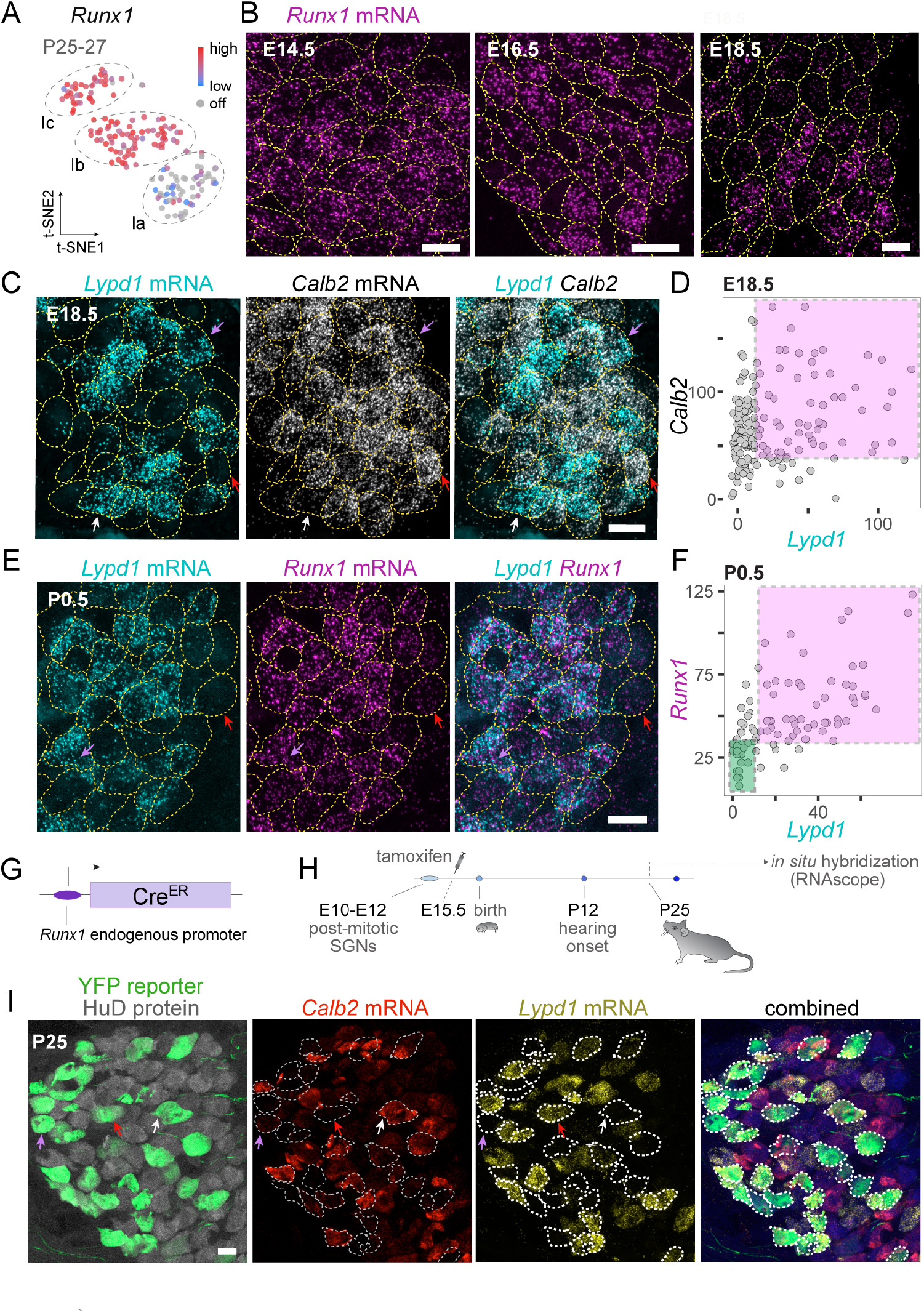
Runx1 expression dynamics in SGNs tracks emergence of subtype identities. (**A**) scRNA-seq of SGNs showed that the transcription factor *Runx1* is expressed selectively in Ib and Ic neurons, suggesting that it could regulate those molecular identities. Plot shows t-SNE embedding of Type I SGN scRNA-seq profiles generated by re-analyzing data in Shrestha et al. (2018). (**B**-**F**) Expression of *Runx1, Calb2*, and *Lypd1* were assessed across development by RNAscope on mid-modiolar sections through the cochlea. **(B)** *Runx1* expression in SGNs is broad at E14.5 (left) and more restricted at E16.5 (middle). By E18.5 (right), cells with high and low levels of *Runx1* can be clearly distinguished. (**C**,**D**) SGN subtype-specific markers such as *Calb2* and *Lypd1* also exhibit restricted expression by E18.5 reminiscent of their expression gradients in adults (reported in Shrestha et al. 2018). Fluorescent puncta counts per SGN is quantified in D. Although the *Calb2* gradient is weak, variability in *Lypd1* expression is strong, with identifiable *Lypd1*^ON^ and *Lypd1*^OFF^ cells (white and red arrows, respectively, C). However, some *Lypd1*^ON^ neurons co-express *Calb2* (purple arrow, D; magenta box, D), suggesting incomplete segregation of these markers at this stage. (**E, F**) At birth, *Runx1* expression remains segregated and co-varies with *Lypd1*, quantified in F. Neurons with high *Runx1* expression tend to be *Lypd1*^ON^ (purple arrows, E; magenta box, F), a known feature of mature Ic SGNs. *Lypd1*^OFF^ neurons express either low or no *Runx1* (red arrows, E; green box, F), which is observed normally in mature Ib and Ia SGNs, respectively. (**G-H**) Strategy for fate-mapping Runx1-positive SGNs. *Runx1*^CreER^ knock-in transgenic mice express the CreER recombinase under regulation of the endogenous *Runx1* promoter. Tamoxifen was injected at E15.5 and SGN fate was assessed at P25 by RNAscope. (**I**) A majority of neurons that had been *Runx1*^+^ embryonically went on to become *Lypd1*^ON^ (Ic, red arrows, I) or *Lypd1*^OFF^ with low *Calb2* expression (Ib, purple arrows, I). A small subset (white arrow, I) acquired high *Calb2* (Ia) identities by P25. Dotted lines around cells in B, C, E, and I represent hand-drawn outlines based on a fluorescent reporter (not shown except in I) in SGN somata expressed genetically (tdTomato driven by the *bhlhe22*^*Cre*^ driver in B, C, E and YFP driven by *Runx1*^*CreER*^ in I). HuD immunostaining in I (gray) shows all SGN somata. Scale bars: 10 microns

### Temporal dynamics of Runx1 expression in SGNs

We surveyed *Runx1* expression in the mouse spiral ganglion from embryonic day E14.5, when differentiating post-mitotic SGNs (Ruben, 1967) are extending peripheral processes towards hair cells (Koundakjian et al., 2007), to E18.5, when SGN processes have reached hair cells (Coate et al., 2015) (E14-E18, Fig. 1B). *Runx1* mRNA transcripts were detected broadly in SGNs *in situ* as early as E14.5 and then became progressively restricted through embryonic development, resulting in clear heterogeneity in *Runx1* levels by E18.5. This stage is also marked by appearance of molecularly distinct SGNs that are OFF for *Lypd1* and express high levels of *Calb2* (Fig. 1C,D), consistent with their complementary expression in mature Ia SGNs (Shrestha et al., 2018). At birth, SGNs that maintain *Runx1* tend to show strong *Lypd1* expression (Fig. 1E,F). Thus, perinatally, *Runx1* expression already overlaps with the Type Ic marker *Lypd1* and is biased toward cells with lower *Calb2* levels. This indicates that spatiotemporal changes in *Runx1* expression coincide with the appearance of molecularly distinct SGNs during late embryonic stages.

To determine the fates of SGN precursors that express *Runx1* embryonically, we used *Runx1*^*CreER*^ mice, which produce CreER under control of the endogenous *Runx1* locus (Fig. 1G,H). *Runx1*^*CreER*^*/+; Ai3/+* mice were administered Tamoxifen at E15.5 to genetically tag Runx1^+^ SGNs with YFP (Fig. 1H). Evaluation of SGN identity by multiplexed fluorescent *in situ* hybridization (RNAscope, Advanced Cell Diagnostics) for *Lypd1* and *Calb2* at P25 revealed that the majority of neurons that were *Runx1*^+^ at E15.5 had taken on Ib (i.e. *Calb2*^MID^ *Lypd1*^OFF^) or Ic (i.e. *Calb2*^LOW^ *Lypd1*^ON^) identities (Fig. 1I). Taken together, these results hint that *Runx1* may be an important player in the generation of molecularly diverse SGNs.

### Effect of Runx1 loss on SGN molecular identity

To directly test whether *Runx1* is necessary for SGN diversification, we abolished *Runx1* function in SGNs by pairing *bhlhe22*^*Cre*^ (Ross et al., 2010), which drives expression of Cre recombinase in SGNs by E9.5 (Appler et al., 2013), with a floxed *Runx1* allele (*Runx1*^*F*^) (Growney et al., 2005) and a fluorescent reporter (*Ai14*). This approach deleted exon 4 of *Runx1*, thereby ablating the Runt domain critical for its DNA- and cofactor-binding abilities (Growney et al., 2005), and simultaneously resulted in tdTomato expression. *Runx1*^*CKO*^ mice are viable and show no obvious behavioral deficits. SGN identities were assessed by conducting single cell RNA-sequencing (scRNA-seq) of neurons from young adult mice (P30-P33) enriched from single cell dissociates of the entire cochlea by fluorescence-activated cell sorting (FACS) (Fig. 2A). Both dimension reduction by Uniform Manifold Approximation and Projection (UMAP) and graph-based clustering of SGN transcriptomic profiles revealed four molecularly distinct subgroups in control animals (Fig. 2B). Three of the subgroups corresponded to Type Ia, Ib, Ic identities and the fourth had a Type II molecular profile. Subtype markers were expressed in expected patterns (Fig. 2C), indicating that this scRNA-seq workflow faithfully captured cell types and gene expression states that represent the major source of molecular heterogeneity among these neurons described previously (Shrestha et al., 2018).

**Figure 2:**
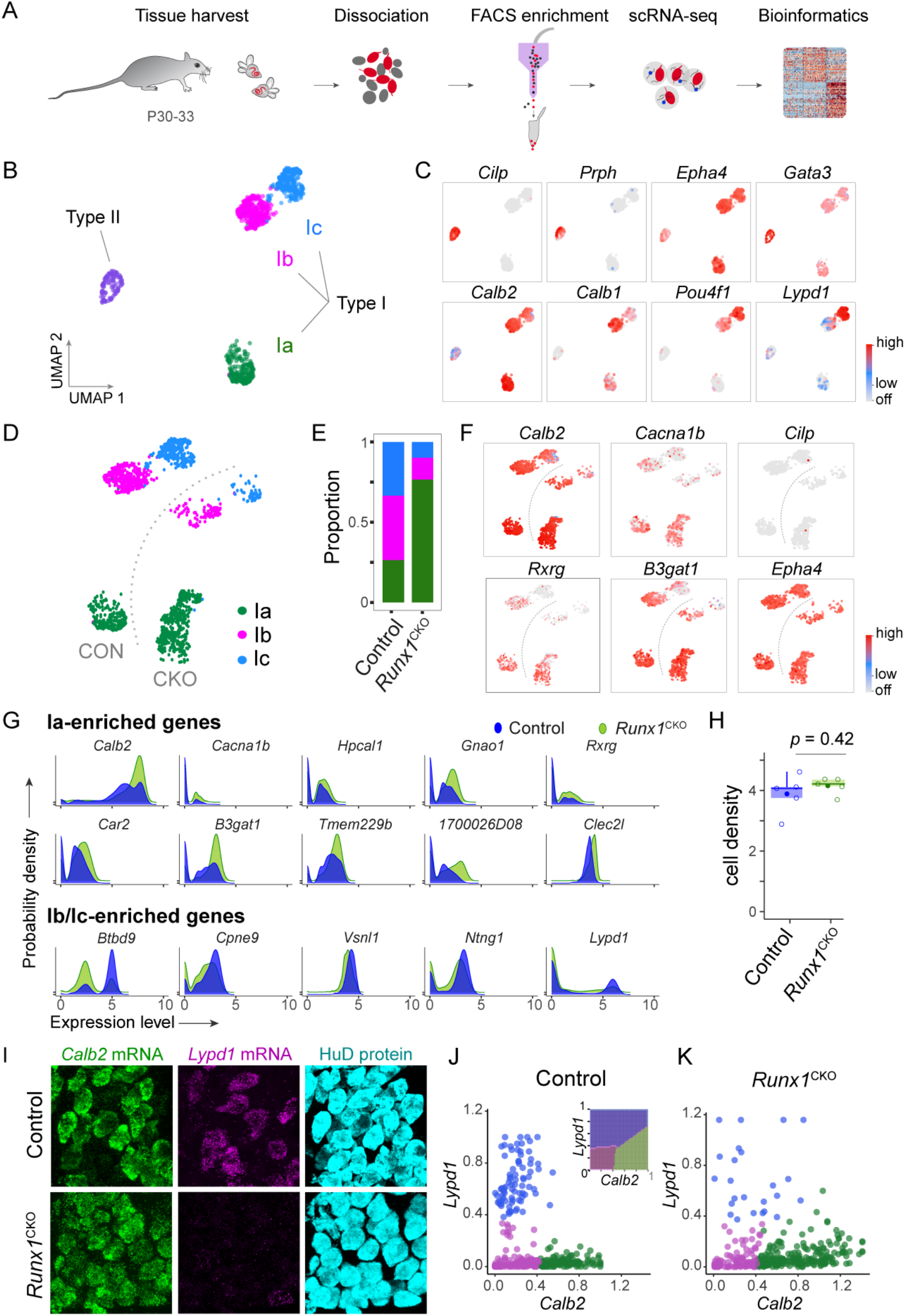
SGN subtype composition is altered upon loss of Runx1 expression. **(A)** Schematic showing the workflow for FACS-based enrichment and subsequent scRNA-seq analysis of SGNs. (**B**) Two-dimensional embedding and unsupervised clustering of neuronal profiles from control animals revealed, as expected, four clusters, three corresponding to Ia, Ib and Ic SGNs and one to Type II SGNs, identifiable based on cluster-specific markers (**C**). (**D**) Type I SGNs from *Runx1*^*CKO*^ mice also segregated into three clusters, shown in a cropped and transformed UMAP plot (see Fig. S1D). (**E**) Neuronal census in each subgroup was drastically different between controls (CON) and mutants (CKO). Ia neurons constituted 26.4% of control Type I SGNs but 76.6% in *Runx1*^*CKO*^ animals. The proportions of both Ib and Ic SGNs were lower in *Runx1*^*CKO*^ (13.6% and 9.7%, respectively) compared to Control (40.2% and 33.4%, respectively). (**F**) The expanded pool of Ia SGNs in *Runx1*^*CKO*^ mice expressed Ia-specific markers (high *Calb2, Rxrg, Cacna1b, B3gat1*) without any change in Type I (*Epha4*) vs. Type II (*Cilp*) markers. Cells remaining in Ib and Ic clusters in the *Runx1*^*CKO*^ group did not acquire Ia markers, indicating that gain of Ia-like transcriptional profiles at the expense of Ib and Ic profiles is the key outcome of Runx1 loss. (**G**) Assessment of gene expression patterns while remaining agnostic to SGN sub-classes revealed that markers known to be Ia-enriched based on previous studies were expressed in more *Runx1*^*CKO*^ mutant cells (green) compared to controls (blue). Conversely, Ib/Ic-enriched genes were underrepresented among *Runx1*^*CKO*^ cells. (**H**) Cell density in the spiral ganglion was statistically identical between the control and *Runx1*^*CKO*^ groups (t-test, *p* = 0.42). (**I**) RNAscope-based evaluation of SGN subtype identities *in situ* revealed flattened *Calb2* gradients and drastic reduction in *Lypd1*^+^ SGNs. HuD is a pan-SGN marker detected by immunohistochemistry. (**J**) K-means clustering of RNAscope-based SGN molecular profiles yielded Calb2^HI^ Lypd1^LOW^ (green), ‘Calb2^LOW^ Lypd1^LOW^ (magenta) and Calb2^LOW^ Lypd1^HI^ (blue) subgroups. A support vector machine (SVM)-based classifier (inset in **J**) was generated using SGN profiles from control animals and used to predict the identities of *Runx1*^*CKO*^ SGNs. (**K**) This analysis revealed severe reduction in the census of Ic-like Calb2^LOW^ Lypd1^HI^ SGN profiles (blue) and expansion of Ia-like Calb2^HI^ Lypd1^LOW^ (green) identities.

scRNA-seq analysis of SGNs from *Runx1*^CKO^ animals revealed a striking change in the distribution of subtype identities. The mutant SGNs segregated into the same four classes, overlapping with those from control animals in UMAP space (Fig. S1C,D). However, although all three Type I subgroups could be detected in both *Runx1*^CKO^ animals and controls, the relative proportions of SGN subtypes was significantly altered: whereas Ia, Ib, Ic neurons were 26.4%, 40.2%, 33.4% of total Type I SGNs in controls, they made up 76.6%, 13.7%, 9.7% in *Runx1*^CKO^ animals, respectively (Fig. S1D, 2D-E). These changes constitute a 90% increase in Ia SGN proportion, and 66% and 71% decreases in those of Ib and Ic SGNs respectively, suggesting that precursors that would normally take on Ib or Ic identities are developing instead into Ia SGNs. This change in distribution cannot be explained by technical variables, as cluster identities did not correlate with measures of single cell library quality and complexity such as number of mRNA and number of genes detected (Fig. S1E,F). Cells in the expanded Ia cluster in the UMAP projection expressed Ia-enriched genes while maintaining differential expression of Type I (*Epha4*) vs. Type II (*Cilp*) SGN markers (Fig. 2F), consistent with the notion that the key change is a switch to Ia identity. Importantly, this interpretation held true even when gene expression trends were analyzed without clustering or projecting onto a low-dimensional space: Ia-enriched genes were expressed at higher levels in more neurons while the opposite was true for Ib/Ic markers in *Runx1*^*CKO*^ SGNs compared to controls (Fig. 2G).

Independent assessment of the SGN phenotype further supports the conclusion that the change in neuronal proportions is due to a conversion of Ib/Ic SGNs into Ia SGNs. Since SGNs undergo cell death during postnatal development and loss of *Runx1* expression is known to affect survival of hindbrain motor neurons (Theriault et al., 2004), we first asked if changes in cell survival can account for altered SGN subtype proportions in *Runx1*^*CKO*^ animals. We found no evidence of difference in neuronal viability upon loss of *Runx1*: neuronal density in the spiral ganglia of *Runx1*^CKO^ animals was statistically indistinguishable from that in controls (Fig. 2H). Next, we assayed gene expression *in situ* by performing RNAscope for the subtype markers *Calb2* and *Lypd1* on cryosections of the cochlea. To interpret these results, K-means clustering was performed on cell-specific gene expression levels determined by quantitative analysis of fluorescent RNAscope puncta using Imaris (Oxford Instruments, UK). SGNs in control animals could be classified into three distinct subgroups (Fig. 2I,J). Consistent with previous reports based on transcriptional profiling (Petitpré et al., 2018; Shrestha et al., 2018; Sun et al., 2018), the subgroups marked by *Calb2*^MID^, *Lypd1*^OFF^ (Ib) and *Calb2*^LOW^, *Lypd1*^ON^ (Ic) comprised nearly 2/3^rd^ of the total SGN population. A support vector machine (SVM)-based classifier built using control data (inset in Fig. 2J, see Methods) grouped *Runx1*^CKO^ SGNs into the same three subgroups, albeit at clearly different proportions—*Calb2*^HI^*Lypd1*^OFF^ Ia SGNs were overrepresented by 77% compared to controls, with marked reduction of *Calb2*^LOW^ *Lypd1*^ON^ Ic identities (Fig. 2K). Taken together, these results support the conclusion that overabundance of Ia SGNs in mice lacking functional *Runx1* in SGNs arises from aberrant apportioning of neuronal molecular identities during development.

### Mixed identities and hierarchical plasticity of gene expression

Neuronal identities can be defined along multiple facets such as connectivity, metabolic signatures, synaptic properties, and intrinsic physiology (Arlotta and Hobert, 2015). Specification of identities can be achieved via transcriptional modulation of all such facets by a master regulator acting through intermediary TFs or by a collection of independent-acting TFs that designate certain neuronal features. Previous work has shown that, among DRG sensory neurons, Runx1 controls non-peptidergic nociceptive neuron identity by regulating both ion channel profiles and innervation patterns within the dorsal horn (Chen et al., 2006). This raises the possibility that Runx1 likewise coordinates multiple aspects of identity in SGNs.

Closer examination of gene expression patterns revealed that Runx1 can have variable effects on SGN subtype identity. By comparison to control Ia, Ib, and Ic SGNs, some *Runx1*^*CKO*^ Ia SGNs seemed to show a complete conversion to the Ia identity: these SGNs expressed many Ia-enriched genes, such as *Calb2, Cacna1b, Rxrg, B3gat1* (Fig. 2F), and also did not express Ib/Ic-enriched genes, such as *Kcnip2, Nalcn, Ncald*, and *Slit2* (Fig. S1G). However, other SGNs in the Ia cluster failed to downregulate all Ib/Ic genes – *Ntng1, Lypd1*, and *Grm8* expression was retained in a subset of *Runx1*^*CKO*^ SGNs that otherwise matched Ia SGNs (Fig. 3B). A broader look at subtype-specific differentially expressed (DE) genes indicated similar trends for more genes, particularly ones that are Ic-enriched (Fig. 3A). Thus, not all genes changed in patterns expected for the change in subtype profiles, raising the possibility that some of the *Runx1*^CKO^ SGNs took on mixed identities.

**Figure 3:**
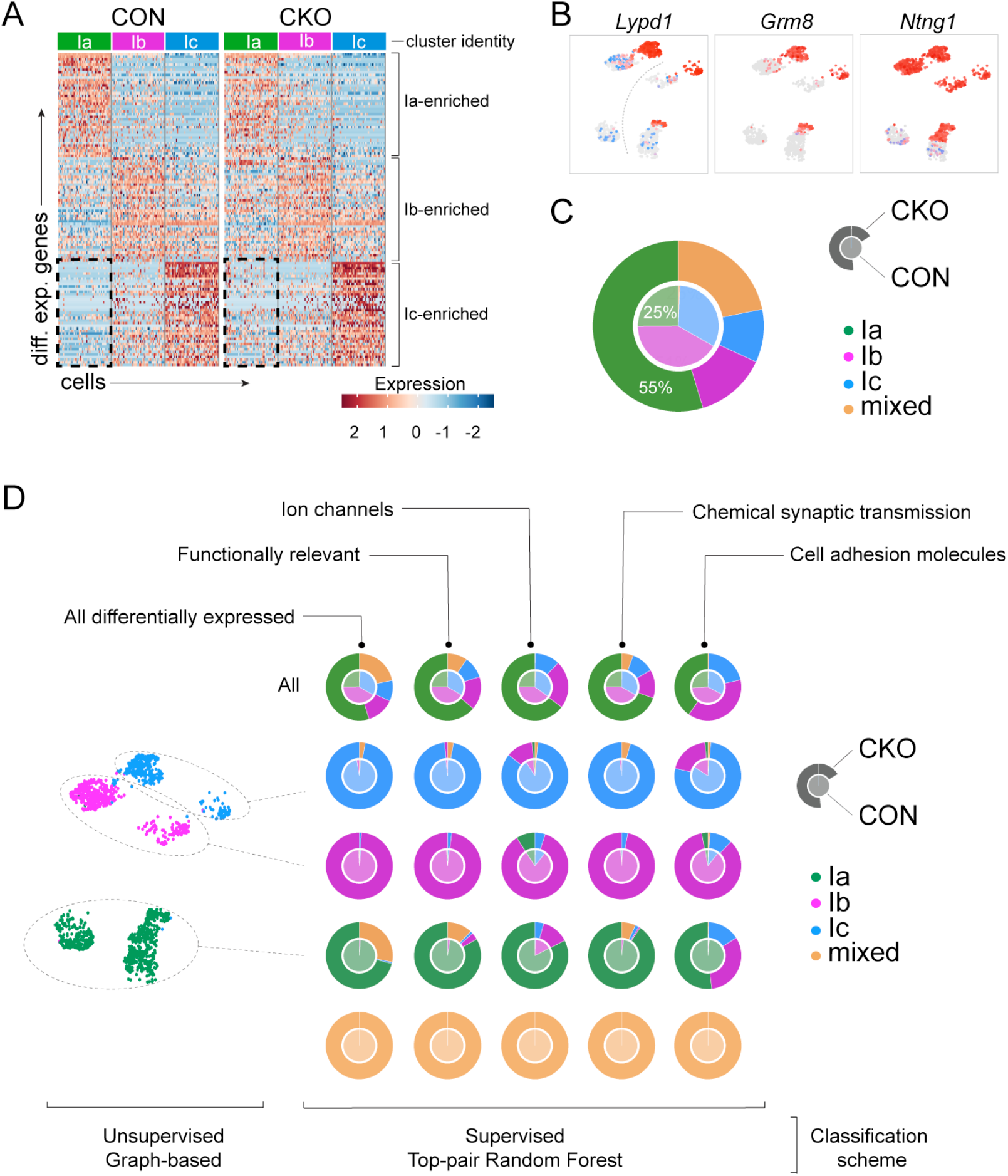
Varying influence of Runx1 loss on different gene categories. (**A**) Heatmap showing top 100 subtype-specific markers for SGNs in control and *Runx1*^*CKO*^ groups. Segregation of subtype markers occurs in *Runx1*^*CKO*^ neurons as it does in Control neurons, but with a key difference: some Ic-specific markers are unexpectedly expressed in Ia neurons in the *Runx1*^*CKO*^ group (compare areas demarcated by dotted rectangles). (**B**) Examples of three Ic-enriched genes expressed ectopically in the *Runx1*^*CKO*^ Ia cluster. (**C**) SGN subtype census determined by a supervised classification scheme designed to detect unresolved or mixed identities. The outer ring represents cell proportions in *Runx1*^*CKO*^ and the center pie represents those in Control. Ia SGNs are over-represented in *Runx1*^*CKO*^ animals by 118% relative to Control, but 22% of SGNs are of mixed identity. (**D**) SGN clusters identified using an unsupervised approach were classified a second time using a supervised approach in order to determine the source of mixed identities and the gene groups contributing to them. Rows correspond to cell subsets while the columns indicate the gene subsets used for supervised classification. The bottom row (yellow) shows how the classifier performed at detecting synthetically created mixed identities spiked into the training dataset. Supervised classification taking all genes differentially expressed among Ia, Ib, Ic Control SGNs (leftmost column) showed that the vast majority of mixed identity SGNs are from the Ia cluster in the UMAP plot. Results of supervised classification using gene subsets drawn from different ontological groups are shown in columns 2-5.

To comprehensively define whether SGNs are cleanly cast into one of three possible molecular identities in *Runx1*^CKO^ animals, as happens in wildtype animals by the 4^th^ postnatal week, we performed supervised clustering of SGN scRNA-seq profiles. Cell identities were assigned based on ensemble learning using DE genes. Additionally, to improve the fidelity of cell class prediction, random gene pairs—as opposed to individual genes—were used to derive a top-pair random forest classifier (TP-RF) (see Methods). We first used the classifier built using all DE genes (n=508) to predict SGN identities in test data from control animals. Considering results of unsupervised clustering as the ground truth, the classifier performed at 98% accuracy overall; only 5% of the 2% error was attributable to incorrectly assigning Ia identity to cells. When applied to *Runx1*^CKO^ SGNs, the classifier yielded predictions in which the proportions of Ia, Ib, Ic SGNs were 51%, 12% and 9%, respectively (Fig. 3C). This represents nearly a doubling of the proportion of Ia SGNs, which normally comprise only ∼25% of the population. In addition, 28% of the mutant SGNs were of mixed identity. Nearly all of these mixed cells were part of the Ia group in a UMAP two-dimensional embedding, confirming earlier hints (Fig. 2F, S1D) that *Runx1*^CKO^ Ia neurons are heterogenous. In fact, although ∼73% of cells in the *Runx1*^*CKO*^ Ia UMAP cluster bore clean Ia identities, the rest were flagged as having mixed identity (Fig. 3D, leftmost column, row 4). Such cases of mixed neuronal profiles are unlikely to be artifacts of our single cell dissociation approach; an algorithmic approach for identifying doublets–joint neuronal profiles of two cells–showed that none could be detected in our scRNA-seq libraries (Fig. S1H-J). Rather, the mixed profiles seem to reflect incomplete conversion to the Ia identity upon *Runx1* loss in a subset of SGNs: adequate number of genes change to classify these neurons as Ia but some Ib and Ic-enriched genes are retained. While these differences could be random, an alternative possibility is that the genes that change expression play a particular role in determining specific neuronal properties.

To determine whether specific facets of SGN identities change upon *Runx1* loss, we built TP-RF classifiers based on more restricted gene sets (Fig. 3D). Predictions of SGN identities were made (Fig. 3D, Table S2) separately using DE genes in the following categories: (1) all genes relevant for neuronal physiology (n=142); (2) genes encoding ion channels (n=67); (3) genes associated with chemical synaptic transmission (n=97); and (4) genes encoding cell adhesion molecules (n=57). Classification based on genes related to neuronal physiology predicted 64%, 16% and 10% Ia, Ib and Ic SGNs, respectively, in the *Runx1*^CKO^ group. We found that 10% of total SGNs had mixed identity, while 83% of SGNs in the Ia UMAP cluster were indeed Ia in supervised classification (Fig. 3D). Classifying based on genes encoding ion channels alone produced a similar result: 65% of total SGNs were Ia, with 82% of cells in the Ia UMAP cluster also predicted to be Ia by the TP-RF classifier (Fig. 3D). Likewise, 70% of total SGNs and 91% of cells in the Ia UMAP cluster were Ia when only genes involved in chemical synaptic transmission were considered (Fig. 3D). In contrast, only 40% of total SGNs were Ia based on cell adhesion molecules in the *Runx1*^CKO^ group. Of the cells in the Ia UMAP cluster, only 52% were assigned Ia identities by the classifier and the remainder were assigned either Ib (32%) or Ic identities (16%) (Fig. 3D). Taken together, although a large majority of SGNs in *Runx1*^CKO^ animals have typical Ia profiles, loss of *Runx1* can yield staggered gene expression outcomes in some SGNs, with more complete switchover of those related to physiological and synaptic function than to connectivity.

### Effects on synaptic location and morphology

In the mature cochlea, most SGNs form a single peripheral synapse whose location along the basal pole of IHC correlates with the neuron’s subtype identity (Liberman, 1982; Shrestha et al., 2018). Ia SGNs make synaptic contacts on the pillar face of IHCs, while Ib and Ic SGNs innervate the modiolar face (Fig. 4A). Additionally, synapses on each side of the IHC exhibit striking differences in morphology and function: the modiolar side tends to have larger presynaptic ribbons (Liberman and Liberman, 2016; Liberman et al., 2011; Shrestha et al., 2018), smaller postsynaptic glutamate receptor patches (Liberman et al., 2011), and active zones where Ca^2+^ influx occur at more depolarized potentials (Ohn et al., 2016)compared to the pillar side. However, it is unclear which of these features may be dictated by SGN subtype identity. Indeed, presynaptic specializations in IHCs are influenced both by IHC-intrinsic polarity (Jean et al., 2019) and by extrinsic signals from SGNs (Sherrill et al., 2019) and olivocochlear efferents (Yin et al., 2014).

**Figure 4:**
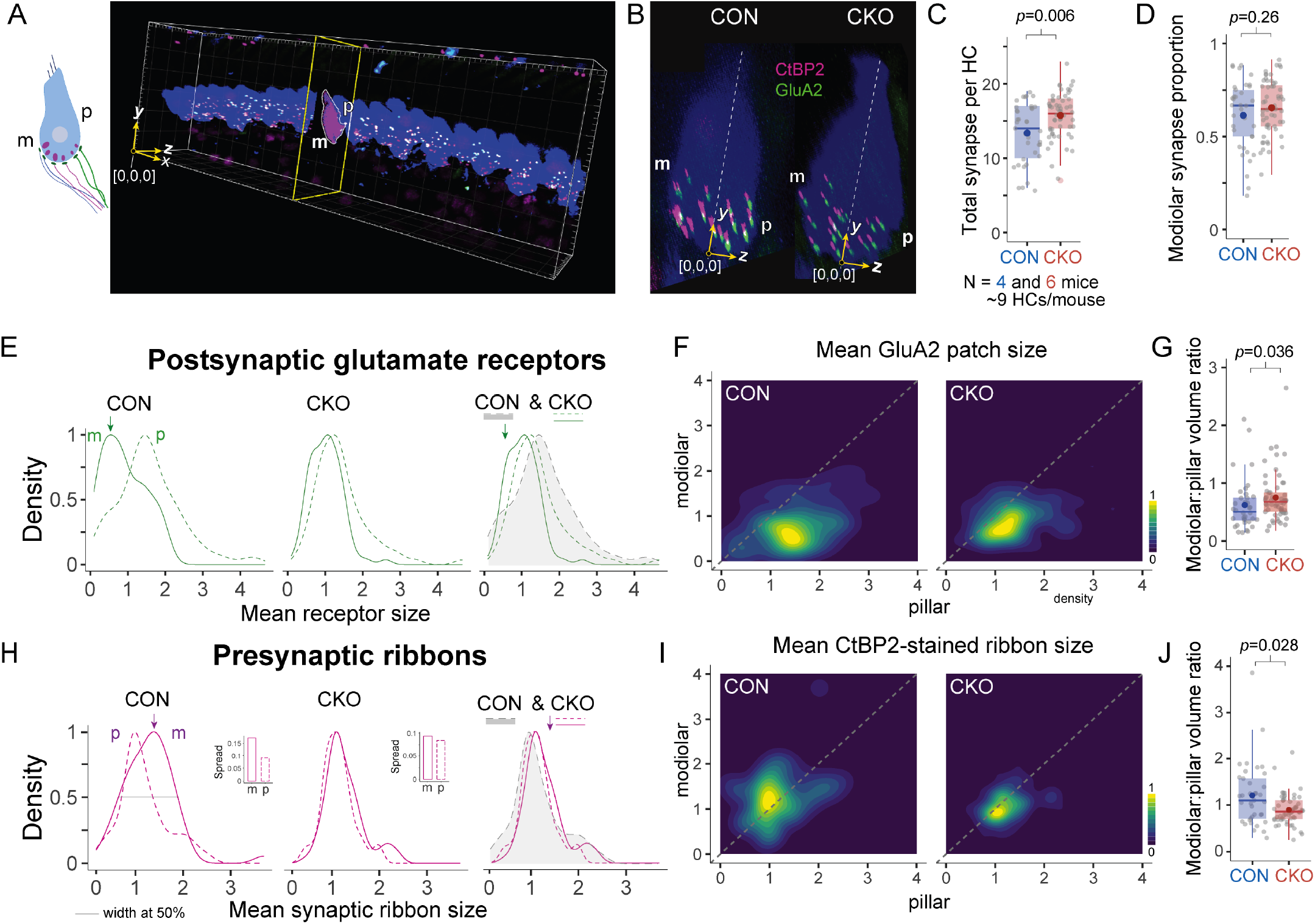
Change and stability in synaptic properties with Runx1 loss. (**A**,**B**) Ia, Ib, and Ic SGN terminals are arranged along the pillar (p) to modiolar (m) axis of the inner hair cell (HC). Synaptic elements marked by CtBP2 (magenta) or GluA2 (green) on the HC surface (blue) were analyzed using Imaris and ImageJ. Synapse location was determined by deriving global image-centric coordinates (yellow axes in A) for each synaptic element using Imaris and determining their position within modiolar or pillar regions defined for each HC after transforming to local HC-centric coordinates (yellow axes in B, see Methods). Images in B are *yz* views of maximum intensity projections along the *x* axis. The projection was clipped to span a single HC. (**C**) A modest increase in synapse number was observed in Runx1^CKO^ mice (15.8±3.0) relative to Control (13.4±3.9) (Wilcoxon Rank Sum Test, *p*=0.006). (**D**) No change was observed in the proportion of synapses on the modiolar face of HCs (Control: 0.61+0.19; Runx1^CKO^: 0.65+0.15; t-test, *p*=0.26). (**E**) Kernel density plots depicting distribution of postsynaptic glutamate receptor (GluA2) volumes, separated by modiolar (m) and pillar (p) location, in Control and *Runx1*^*CKO*^ groups. The rightmost plot has an overlay of three distributions from those to the left. Arrows mark the peak of the distribution for modiolar synapses in CON to highlight rightward shift in the CKO group. (**F**) Two-dimensional density plots of mean GluA2 volumes on modiolar vs. pillar sides of each HC. Dotted line depicts line of identity corresponding to equal sizes on modiolar and pillar sides. (**G**) Comparison of mean modiolar:pillar GluA2 volume ratio of each HC. Mean ratio in the Control group was well below 1 (0.62±0.08) and the ratio in the *Runx1*^*CKO*^ group was significantly higher (0.75±0.06), indicating weakening of the size gradient (Wilcoxon Rank Sum Test, *p*=0.036). (**H**) Distribution of presynaptic ribbon volumes in Control and *Runx1*^*CKO*^ groups. Column charts in inset compare the width of the density curves at 0.5 probability density (gray line) as a measure of spread in size distribution between modiolar (m) and pillar (p) regions. Arrows indicate peak of modiolar size distribution in CON to highlight leftward shift in the CKO group. (**I**) Two-dimensional density plots of mean ribbon volumes on modiolar vs. pillar sides of each HC. Dotted line depicts a line of unity corresponding to equal sizes on modiolar and pillar sides. (**J**) Ratios of mean presynaptic ribbon volumes on modiolar vs. pillar sides of each HC were compared. Mean ratio in the Control group was above 1 (1.21±0.11), indicating a modiolar-to-pillar gradient. The ratio in the *Runx1*^*CKO*^ group was significantly lower and below 1 (0.90±0.04), indicating loss of the size gradient (Wilcoxon Rank Sum Test, *p*=0.028). C,D,G,J are standard box-and-whisker plots with added colored dots denoting groupwise means.

We first asked whether loss of *Runx1* function results in shifts in synaptic position that match the change in proportions of Ia, Ib, and Ic SGNs revealed by scRNA-seq profiles (Fig. 2). Despite drastic reductions in Ib/Ic identities, the overall number of synapses onto IHCs was not decreased in *Runx1*^CKO^ animals compared to controls (Fig. 4A, B, S3A; see Methods). In fact, a modest increase in total synapse number per IHC was observed compared to controls (Fig. 4C, Control:13.4±3.9; Runx1^CKO^:15.8±3.0; Wilcoxon Rank Sum Test, *p*=0.006). Additionally, even though Ia SGNs were gained at the expense of Ib and Ic SGNs, the proportion of synapses on the modiolar side in *Runx1*^CKO^ mice remained unchanged (Fig. 4B,D), indicating that synapse location was unaffected (Control: 0.61+0.19; Runx1^CKO^: 0.65+0.15; t-test, *p*=0.26).

To determine whether synaptic morphology tracks with SGN identity independent of location, we analyzed the volumes of glutamate receptor (GluA2) densities on each side of the IHC. In control animals, mean GluA2 volumes for individual IHCs showed a clear trend toward larger sizes on the pillar compared to the modiolar side (Fig. 4E) consistent with previous reports (Liberman and Liberman, 2016; Liberman et al., 2011). Upon loss of *Runx1* expression in SGNs, this difference disappeared, owing to a shift toward larger GluA2 volumes on the modiolar side (Fig. 4E,F). Two-dimensional representations of modiolar and pillar GluA2 volumes for individual IHCs revealed a distribution below the line of identity in scatter (Fig. S3B) and density plots (Fig. 4F) for control animals, indicative of a pillar-modiolar gradient. However, in *Runx1*^CKO^ animals, mean GluA2 volumes shifted closer to the line of identity, indicating weakening of the gradient. This held true even when we measured the GluA2 gradient by calculating the modiolar:pillar volume ratio on a HC-by-HC basis: the mean ratio was well below 0.62±0.08 in control animals but increased to 0.75±0.06 in *Runx1*^CKO^ animals (Wilcoxon Rank Sum Test, *p*=0.036; Fig. 4G). Thus, *Runx1* mutant SGN terminals on the modiolar side exhibited features normally associated with Ia SGN terminals on the pillar side.

Since previous studies have reported opposing gradients of postsynaptic GluA2 and presynaptic ribbon volumes at IHC-SGN synapses (Liberman and Liberman, 2016), we wondered whether presynaptic changes follow shifts in GluA2 size distribution. Importantly, IHCs do not express *Runx1* (Kolla et al., 2020; Scheffer et al., 2015)and are not targeted by the *bhlhe22*^Cre^ driver line (Meng et al., 2019), so direct genetic perturbation in *Runx1*^CKO^ animals is restricted to SGNs in the cochlea. Presynaptic ribbon sizes, measured as mean volumes of anti-CtBP2 stained puncta, tended to be larger on the modiolar side than those on the pillar side in control animals (Fig. 4H), consistent with previous reports (Liberman and Liberman, 2016; Shrestha et al., 2018). This difference disappeared completely upon loss of *Runx1* expression in SGNs (Fig. 4H,I); both pillar and modiolar ribbon volumes in *Runx1*^CKO^ animals matched the size distribution of pillar-localized ribbons from control animals (Fig. 4H). Two-dimensional scatter (Fig. S3C) and density plots (Fig. 4I) of modiolar vs. pillar ribbon volumes for individual IHCs revealed a distribution above the line of identity for control animals, indicative of a modiolar-pillar gradient. However, in *Runx1*^CKO^ animals, the ribbon volume distribution lay below the line of identity, indicating collapse of the gradient. This interpretation was also supported by measurements of the modiolar:pillar volume ratio per IHC: the mean modiolar:pillar ratio was 1.21±0.11 in control animals but significantly lower (0.90±0.04) in *Runx1*^CKO^ animals (Wilcoxon Rank Sum Test, *p*=0.028, Fig. 4J). Furthermore, we noted a distinctly larger IHC-to-IHC variation in mean ribbon sizes on the modiolar as compared to the pillar side in the control group (see gray line and inset in Fig. 4H). This suggests that the extent of morphological heterogeneity across HCs is a distinguishing feature of these two presynaptic domains. This variation in the volume of modiolar ribbons was lost in *Runx1*^CKO^ animals, with more homogeneous size distribution akin to those on the pillar side (inset in Fig. 4H).

Thus, upon deletion of *Runx1*, both pre- and post-synaptic elements of IHC-SGN synapses changed in such a way that size gradients collapsed and modiolar synapses were now morphologically indistinguishable from pillar synapses. In addition, all the changes occurred in a direction that produced a final outcome matching the morphology of pillar synapses in control animals. Taken together, a broad shift toward Ia molecular identity in *Runx1*^CKO^ animals was accompanied by congruent changes in pre- and post-synaptic components toward Ia-specific features, even as synaptic location remained unaltered.

### Neural response with altered SGN subtype census

Our studies suggest that there is an expansion of neurons with Ia-specific (high-SR like) properties in *Runx1*^CKO^ animals, notably based on expression of functionally relevant genes. Therefore, we set out to characterize SGN function in these animals by performing auditory brainstem response (ABR) recordings. Peak 1 (P1, arrow in Fig. 5A) in an ABR waveform corresponds to the composite neural activity of all SGNs (Melcher and Kiang, 1996). Since high-SR fibers have relatively larger and more synchronized onset responses than low-SR fibers (Bourien et al., 2014), they contribute more to ensemble neural potentials like the ABR. Comparison of ABR P1 amplitude thus offers a window into changes in SGN subtype composition. For instance, loss of low-SR SGN connectivity peripherally due to noise exposure reduces suprathreshold P1 amplitudes without affecting ABR threshold (Furman et al., 2013; Kujawa and Liberman, 2009).

**Figure 5:**
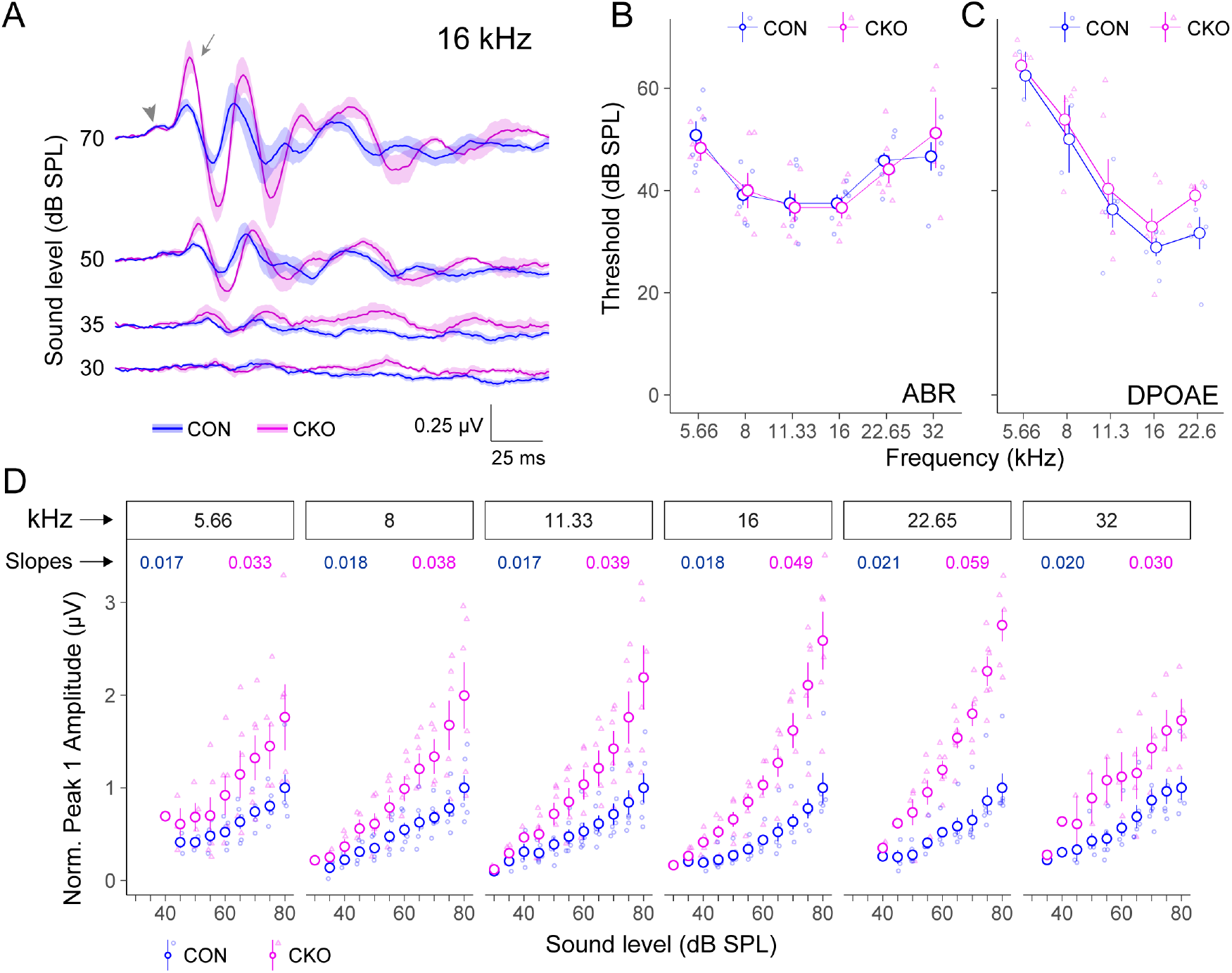
Altered neural response to sound in the cochlea of *Runx1*^*CKO*^ mice. Auditory brainstem response (ABR) and distortion product otoacoustic emission (DPOAE) were recorded from Control (blue) and *Runx1*^*CKO*^ (magenta) mice. (**A**) Averaged waveforms of ABRs recorded upon presentation of 16 kHz tone burst at varying sound levels. While SGN responses in peak 1 (arrow) grow as expected with increase in sound level in Control animals, the growth is much larger in the *Runx1*^*CKO*^ group. Arrowhead indicates summating potential. (**B**,**C**) No difference in ABR (B) or DPOAE (C) thresholds was observed across all sound frequencies tested (Table S4). (**D**) Normalized peak 1 amplitude vs. stimulus level plotted for each sound frequency. Slopes of the input-output growth functions for the *Runx1*^*CKO*^ group were larger by a factor of 1.5 or more compared to the Control group. Plots in B and C share the same y-axis labels. All plots in D share the same y-axis. All plots show mean ± SEM for the respective groups. Results of statistical tests for data in B,C,D can be found in Table S4.

ABR recordings (Fig. 5A, S4) revealed normal thresholds in *Runx1*^CKO^ mice compared to controls when tested with 5.66, 8, 11.33, 16, 22.65 and 32 kHz pure-tone bursts (Fig. 5B, Table S4). However, P1 amplitudes at suprathreshold sound pressure levels were larger, reflecting enhanced neural responses, at level-matched stimulus presentations in *Runx1*^CKO^ animals (Fig. 5A,D, Table S4). This translated to steeper growth functions in *Runx1*^CKO^ compared to littermate controls, with slope increases ranging from 1.5 to 2.7-fold (Fig. 5D). Differences were observed across the tonotopic axis, but the strongest effect was observed in mid-cochlear regions. Notably, indicators of hair cell function were unaffected, with no detectable change in DPOAE threshold (Fig. 5C) or summating potential (arrowhead, Fig. 5A). Thus, the observed change in auditory nerve responses is likely due to altered SGN properties. These data show that the observed molecular changes in gene expression have functional consequences, with an expansion of SGNs with Ia identity resulting in ABR responses predicted for a cochlea with more high-SR SGNs.

### Change in SGN identity upon postnatal loss of Runx1

Although molecularly distinct immature SGNs are present by late embryogenesis, perturbation of pre-hearing activity can redirect Ib and Ic SGNs towards the Ia fate (Shrestha et al., 2018; Sun et al., 2018). Since a similar phenotype occurs in *Runx1*^CKO^ animals, we wondered whether *Runx1* also influences this latent capability for postnatal plasticity in SGNs.

Fate-mapping approaches support the idea that SGN subtype identity remains flexible shortly after birth. *Runx1*^*CreER*^*/; Ai14/+* mice were given Tamoxifen at P1-P3, and then SGNs were harvested at P30 and profiled by scRNA-seq (Fig. 6A). SGN transcriptional identities are distinct before birth, with *Runx1* expression already restricted to Ib and Ic SGN precursors (Petitpré et al., 2022). Thus, our fate-mapping strategy should tag neurons that have begun to take on Ib/Ic-specific molecular profiles. We found that while many neonatal *Runx1*+ SGN precursors do indeed acquire a Ib/Ic identity, Ia identities are also possible. Unsupervised clustering analysis revealed that SGNs that expressed *Runx1* at P1-P3 (and are therefore tdTomato+) can become any of the 3 subtypes of Type I SGNs (Fig. 6B). However, nearly 80% of *Runx1*-expressing SGNs retained their tentative Ib and Ic identities while the rest switched to Ia identity (Fig. 6D). Thus, although SGNs are molecularly distinct at P3, these identities can still change, consistent with previous reports that cochlear activity influences SGN subtype proportions (Shrestha et al., 2018; Sun et al., 2018).

**Figure 6:**
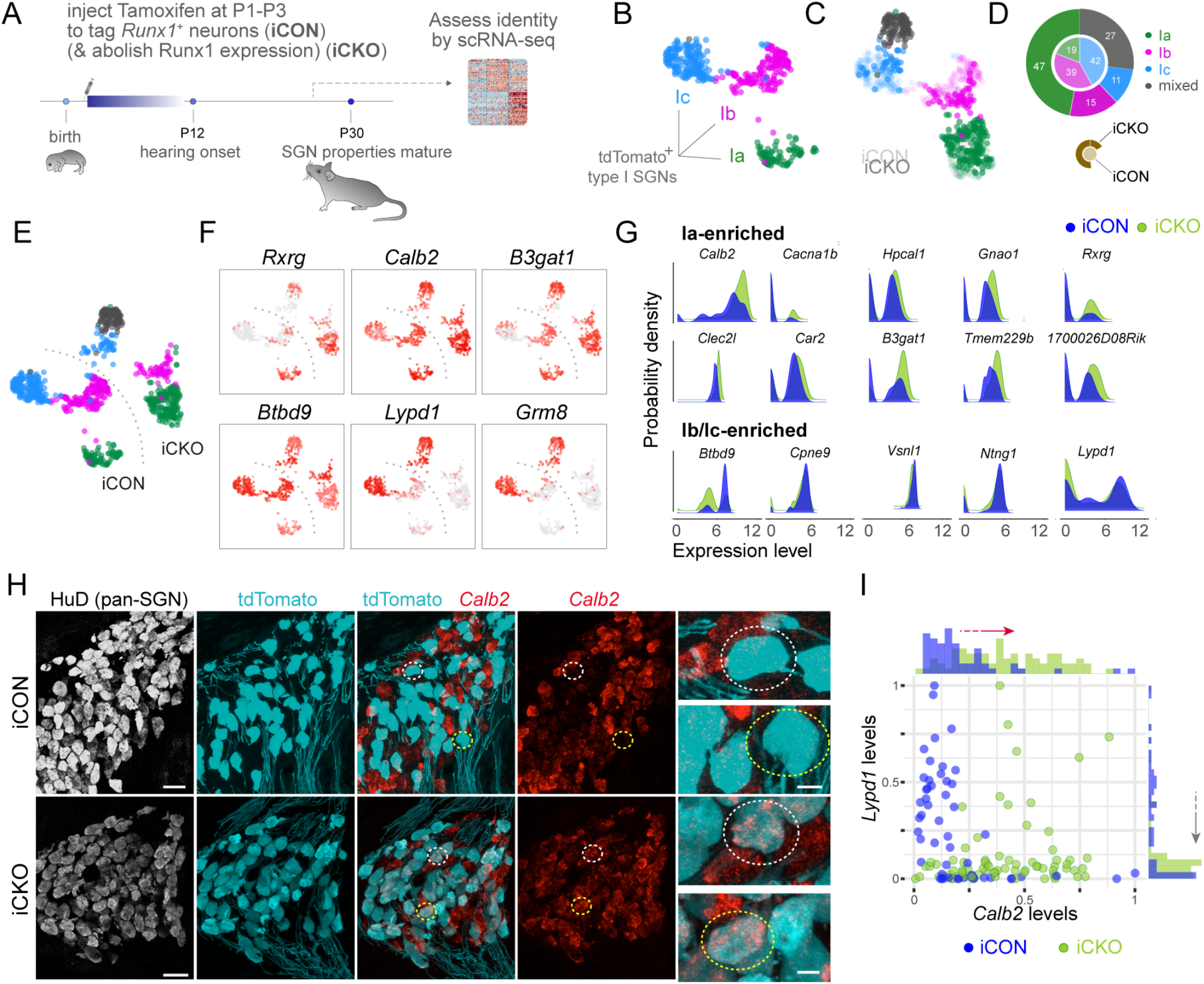
Postnatal Runx1 loss redirects SGN subtype identities. (**A**) Schematic depicting the scRNA-seq workflow for fate-mapping of Runx1^+^ neurons with (iCKO) or without (iCON) postnatal loss of Runx1 expression. (**B**) UMAP embedding of scRNA-seq profiles of SGNs from the iCON group revealed three clusters corresponding to Ia, Ib, Ic subtypes, with a large majority of neurons located in the Ib and Ic clusters. (**C**) UMAP embedding of scRNA-seq profiles of SGNs from the iCON (fainter colors) and iCKO (brighter colors) groups revealed that clusters for the two groups largely overlap but differ in terms of cell census in each cluster. A subset of iCKO cells adjacent to the Ic cluster stands out as a distinct fourth cluster (grey). (**D**) Proportions of SGN subtypes predicted by a supervised classification approach structured to discern Ia, Ib, Ic or mixed identities. Loss of Runx1 postnatally resulted in >2.5 fold increase in the proportion of cells acquiring Ia identity at the expense of both Ib and Ic identities. In addition, more than a quarter of the SGNs are of mixed identity. Numbers indicate % of each SGN subtype within a group. (**E**) UMAP embeddings in which SGNs from the two groups and the clusters they fall in are identified. (**F**) Feature plots of various subtype markers revealed that the expanded pool of Ia SGNs in iCKO express Ia- and turn off Ic-specific genes. The fourth cluster (grey in E) contains cells that co-express Ia- and Ic-specific genes. (**G**) Analysis of gene expression patterns without first classifying SGNs indicate increased representation in the overall population of gene signatures associated with the Ia identity. (**H, I**) Changes in SGN gene expression patterns were also analyzed *in situ* by RNAscope to mid-modiolar sections of the P26 cochlea, along with immunostaining for HuD (white) and tdTomato (blue) to visualize SGNs. Overall, the *Calb2* gradient (red) was shallower among SGNs of iCKO mice (bottom) compared to those in the iCON group (top). Dotted circles highlight tdTomato+ neurons with low *Calb2* level in the iCON group but high *Calb2* level in the iCKO group. Quantification is shown in a scatterplot of *Calb2* and *Lypd1* levels in fate-mapped tdTomato+ SGNs from iCON (blue) and iCKO (green) animals (I). Histograms depict changed distributions of *Calb2* (top) and *Lypd1* (right) expression levels, respectively. Most iCON SGNs are Lypd1^HI^ Calb2^LOW^ or Lypd1^OFF^ Calb2^LOW^ (i.e., Ic- or Ib-like, respectively). In contrast, those from iCKO animals are Lypd1^OFF^ Calb2^LOW^ or Lypd1^OFF^ Calb2^HI^ (i.e., Ib- or Ia-like, respectively). In addition, a subset of cells is Lypd1^HI^ Calb2^HI^ (i.e., of mixed identity). These trends also hold true at the level of individual genes: histograms show shifts toward higher *Calb2* expression (red arrow, top) and lower *Lypd1* expression (gray arrow, right) among fate-mapped SGNs upon postnatal loss of Runx1 expression.

These findings raised the possibility that maintenance of Runx1 is a key step in the consolidation of Ib/Ic SGN identity. To test this idea, we ablated *Runx1* postnatally (P1-P3) in *Runx1*^CreER^/*Runx1*^F^; *Ai14/+* animals (“*Runx1*^iCKO^”). Indeed, this perturbation fundamentally changed the fate outcome of SGNs: despite having intact *Runx1* expression up to P3—and being destined to retain Ib/Ic identity with ∼80% probability as described above—nearly half of the *Runx1*^iCKO^ SGNs shed their Ib/Ic identities and instead become Ia SGNs, as assessed by scRNA-seq of tdTomato+ SGNs (Fig. 6C,D). This represents nearly 150% increase in the probability of becoming Ia SGN compared to controls. The decrease in probabilities of retaining Ib and Ic identities are 62% and 74%, respectively. Notably, 27% of *Runx1*^iCKO^ SGNs acquire ‘mixed’ identities. Closer examination of genes expressed in this subgroup of mutant SGNs revealed that the mixture of identities is a result of failure to shed Ic gene signatures despite having acquired Ia gene profiles (Fig. 6E,F). For example, *Runx1*^iCKO^ SGNs expressed *Rxrg, Calb2, BtBd9* and *B3gat1* at levels typical of control Ia SGNs, but failed to turn off the Ic-specific markers *Lypd1* and *Grm8*. These differences are unlikely to be driven by clustering-associated artifacts because examination of population-wide gene expression trends without any regard to cell identity also revealed increased representation of Ia-associated genes and decreased expression of Ib/Ic-associated genes (Fig. 6G). These findings show that Runx1 also acts postnatally to steer differentiating SGNs away from the Ia identity.

Consistent with scRNA-seq results, similar changes in SGN identity were observed using multiplexed fluorescent *in situ* hybridization (RNAscope) in tissue sections of the cochlea. Most neurons that were tagged with tdTomato—indicating that they expressed *Runx1* before P3—went on to become *Calb2*^LOW^ Lypd1^ON^ (Ic) or *Calb2*^MID^Lypd1^OFF^ neurons (Ib) in mature animals (Fig, 6H,I). In contrast, when we ablated *Runx1* expression shortly after birth, we observed fewer tdTomato+ cells expressing *Lypd1* and more with higher *Calb2* levels (Fig. 6I). Indeed, tdTomato+ SGNs frequently expressed high *Calb2* levels and a subset even co-expressed *Calb2* and *Lypd1*; both of these expression patterns were rare in the control group (Fig. 6H,I). Thus, SGN identities are malleable postnatally and maintenance of Runx1 is required for final consolidation of subtype identity.

## Discussion

Across sensory modalities throughout the animal kingdom, sensory neurons with heterogeneous functional and connectivity profiles are used to encode complex stimuli. In contrast to the stark molecular differences that distinguish fundamentally different neuron types, such as excitatory and inhibitory neurons, molecular variation within a sensory neuron population is subtler and is layered on top of shared programs of gene expression. This raises an important question in neural development: how are functionally important differences within neurons of a single modality established? Here, we show that sensory neurons in the cochlea, the spiral ganglion neurons (SGNs), acquire subtype-specific properties through flexible use of an intrinsic transcriptional template. The key player is the transcription factor Runx1, which coordinates broad transcriptional changes such that SGNs take on Ib or Ic identities rather than Ia identities. These molecular changes were accompanied by predicted changes in synaptic heterogeneity and in physiological responses, but not in synaptic position. Although loss of Runx1 often led to a complete identity switch, some SGNs showed stronger changes in subtype-specific genes related to neuronal function than those that encode cell adhesion molecules, indicating that identity-related genes may be controlled hierarchically. Consistent with the fact that changes in cochlear activity can alter SGN subtype proportions, Runx1+ progenitors are not fully committed to the Ib/Ic identity and switch to Ia identity upon loss of Runx1 just after birth. Thus, the final proportion of SGN subtypes may be shaped by an intersection between activity and Runx1’s intrinsic ability to endow Ib and Ic SGNs with properties needed for encoding sound.

Our data support the idea that Runx1 is a molecular switch that promotes Ib and Ic identities while simultaneously repressing Ia identities. As assessed by both supervised and unsupervised analysis of gene expression, embryonic loss of *Runx1* creates an excess of Ia SGNs, a result that would not be possible if those cells simply lost expression of Ib/Ic-enriched genes without gaining Ia-specific molecular signatures. One interpretation is that Runx1 directly activates genes associated with Ib/Ic identities and directly represses those that confer Ia identity. Such a capacity to orchestrate cellular identity via both positive and negative regulation of gene expression is not unprecedented for *Runx1* (Chen et al., 2006). It is also possible that Runx1 acts indirectly through expression of an as-yet unidentified intermediate TF that controls Ia identity. How distinctions between Runx1-dependent Ib and Ic SGN identities emerge remains to be determined.

Growing evidence indicates that neuronal identity is established in a sequential and modular fashion that includes early acting master regulators, intermediate switches such as Runx1, and late acting terminal selectors, which are transcription factors that induce and maintain specific batteries of genes needed for mature function (Hobert, 2011, 2016). Runx1’s effects on SGN diversification may be carried out through the action of terminal selector-like transcription factors that dictate different subtype-specific features. For instance, Mafb seems to control postsynaptic differentiation without affecting intrinsic firing properties (Yu et al., 2013), whereas Pou4f1 is important for different SGN subtypes to instruct appropriate pre-synaptic differentiation in the IHC (Sherrill et al., 2019). Our characterization of pre- and post-synaptic properties, synapse location as well as neural response to sound upon loss of *Runx1* serves as a valuable platform upon which the TF regulatory logic of SGN diversification can be further studied.

Although SGN identity and auditory function were dramatically altered in *Runx1*^CKO^ mice, some SGNs were only partially converted, suggesting that Runx1’s effects on gene expression are influenced by other factors. With both embryonic and postnatal deletion of *Runx1*, many cells seemingly shed their Ib/Ic identities—and became clean Ia’s —even as others held on to parts of them. Why Ib/Ic gene retention occurs only in a subset of cells is not clear. While some of the clean Ia’s may never have expressed *Runx1*, this would only explain the presence of mixed identities after embryonic deletion; for the postnatal studies, we assayed only those SGNs that were already following the Ib/Ic trajectory. Whether cells with mixed identities represent a subgroup that are inherently less plastic than others and how they may be distributed along the cochleotopic axis, which contains a gradient of cellular maturation, are intriguing questions for future studies. One possibility is that the cells with mixed identities were in a more differentiated state at the time of *Runx1* loss than those that fully converted. It is also possible that activity-dependent processes impact sensitivity to Runx1. Regardless of how Runx1 activity is modulated, our data suggest that there is a hierarchy of biological features with some locked in place early while others remain amenable to change. The rigidity in expression of cell adhesion molecule-associated genes implies that SGN connectivity is far more stable than functional properties through early development. Consistent with this idea, synaptic location along the basal pole of IHCs remained unaltered despite an overabundance of Ia SGNs in *Runx1*^*CKO*^ animals. The implications of such a heterogeneous transcriptional regulatory landscape, featuring rigidity and flexibility in expression across different gene families, remain to be elucidated.

The discovery that SGNs can change their identity in the absence of obvious changes in synapse number or position indicates that mechanisms other than circuit rewiring contribute to developmental plasticity and underscores the idea that stable circuits with flexible physiology are also part of the repertoire of neural development. Throughout the nervous system, neural connections established early in life are subject to change: some are pruned while others are consolidated to yield mature neural circuits. Anatomical refinement of juvenile circuits depends critically on spontaneous or sensory input-driven neural activity (Faust et al., 2021; Katz and Shatz, 1996; Kirkby et al., 2013; Leighton and Lohmann, 2016). In addition to influencing the pattern of synaptic connections, activity can also alter the functional architecture of neural circuits by shaping cell identities (Cheng et al., 2022; Shrestha et al., 2018). Our data further demonstrate that molecularly distinct precursors can switch developmental trajectories postnatally and introduce Runx1 as a mediator of this switch. In the somatosensory system, developing primary neurons also go through transient stages and then commit to specific subtype identities by maintaining expression of key transcription factors in response to target-derived cues (Sharma et al., 2020). Thus, activity-induced changes in functional identity may be a broadly active mechanism used to fine-tune circuit organization, perhaps akin to the use of target-dependent neuronal survival to establish the basic pattern of connectivity during initial wiring (Kuno, 1990). For sensory systems, such a mechanism can be used to adjust subtype proportions based on the needs of the animal. Indeed, SGN subtype proportions differ along the tonotopic axis in mice and there is a higher proportion of Ia SGNs in the apex of the cochlea in gerbils (Bourien et al., 2014).

Given their disproportionately large number of Ia SGNs, *Runx1*^CKO^ animals are a valuable model for understanding the significance of SGN diversity for sound encoding in the auditory nerve. Although significant insights have been gained from animal models with depletion of Ic SGNs due to noise exposure or old age, *Runx1*^CKO^ animals offer the opportunity to investigate how stimulus coding changes when the neurons are predominantly of one functional class without reducing the total neuron number, without changing the overall circuit organization, and outside the context of an injury model. The discovery that responses to suprathreshold stimuli are larger and grow faster in *Runx1*^CKO^ animals emphasizes the fact that most SGNs in these animals are functionally behaving like low-threshold Ia neurons. Thus, further analysis of these animals can reveal how such a change in neuronal composition impacts the animal’s perception of sound. Since loss of Ic SGNs is predicted to make it harder to hear in noisy environments, the postnatal malleability of SGN identity raises the possibility of using Runx1 to restore SGN diversity and hence mitigate the effects of age-related hearing loss.

## Data and code availability

Raw scRNA-seq data generated in this study, processed gene expression matrices, and related metadata have been deposited at the NCBI Gene Expression Omnibus (GEO) repository with accession numbers GSE23193258 and GSE210216. Custom R scripts are available upon reasonable request.

## Acknowledgements

This work was supported by NIH/NIDCD grants R01DC009223 to L.V.G. and R21DC018356 to B.R.S., and a Lefler Postdoctoral Fellowship at Harvard Medical School to B.R.S. We thank Dr. Bernardo Sabatini (Harvard) for use of the 10x Chromium Controller, Dr. M. Charles Liberman (Mass Eye and Ear, Harvard) for helpful discussions, Daniela Gutierrez (Harvard) for assistance with cell sorting at the HMS Immunology Flow Cytometry Core Facility, and Drs. Gord Fishell (Harvard), Qiufu Ma (Boston Children’s, Harvard) and M. Charles Liberman for critical feedback on the manuscript.

## Author Contributions

B.R.S. conceived and supervised the project, designed the experiments, performed scRNA-seq, analyzed the data, secured funding, and prepared the manuscript. L.W. performed all experiments except scRNA-seq, analyzed the data, and prepared the manuscript. L.V.G. conceived and supervised the project, secured funding, and prepared the manuscript.

## Declaration of interests

The authors declare no competing interests.

## STAR Methods

### Histology, immunohistochemistry and *in situ* gene expression analysis

Embryonic heads were promptly collected after euthanasia of the pregnant dam by CO_2_ overexposure and drop-fixed in 4% paraformaldehyde (PFA) in 1x PBS. Animals older than P21 were anesthetized via isoflurane exposure by the open-drop method and perfused with 4% PFA in 1x PBS. Both cochleae were dissected out from the temporal bones and cochleostomy was performed using the tip of a scalpel blade at the basal turn to facilitate PFA entry into the cochlea. For tissues used for *in situ* hybridization assays, post-fixation was performed at 4°C for 2 hr before decalcification in 120 mM EDTA in 1xPBS for 48-72 hr at 4°C. They were then equilibrated in increasing sucrose concentrations at 4°C before cryopreservation and sectioning. For tissues used for synapse staining in whole mount preparations, post-fixation was performed for 1 hr at RT, followed by decalcification in 120 mM EDTA in 1xPBS for 48-72 hrs at 4°C. Cochlea were then micro-dissected into apical, middle, and basal turns. Following sucrose permeabilization with a freeze-thaw cycle in 30% m/v sucrose in 1xPBS, non-specific expression was blocked with blocking solution (5% v/v normal donkey serum, 5% v/v normal goat serum, and mouse fab fragments) for 1 hr. Tissues were permeabilized with 1% Triton-X in 1xPBS (PBST). Cochlear turns were stained overnight with anti-Pvg, anti-CtBP2, anti-GluA2, and anti-Nfl and subsequently in appropriate secondary antibodies, both at 37°C. Rinses after primary and secondary antibody incubations were done using 1% PBST every 15 mins for 1 hr at RT.

Anti-HuD staining for assessing SGN density was done using a standard immunohistochemistry protocol as described previously (Shrestha et al, 2018) but with the following addition: at the start of the protocol, sections were post-fixed in 4% PFA for 10 min, rinsed with 0.02% PBST, treated with Type III protease (ACD) for 10 min, rinsed in 0.02% PBST for 5 min, then blocked in consecutive 30 min segments each using 5% normal donkey serum and Fab fragments before proceeding with primary antibody incubation.

For mRNA detection by RNAscope (Advanced Cell Diagnostics), the manufacturer’s protocol was used with the exception that at the end of the protocol, tissues were stained overnight with anti-HuD to mark cell bodies and incubated in the appropriate secondary antibodies for 2 hr at RT the next day. In some cases, tissues were also stained overnight with anti-dsRed to amplify endogenous tdTomato signal.

### Transcriptomic profiling

SGNs were collected from P30-P33 mice of both sexes with the following genotypes: 1) Control: *bhlhe22*^*Cre*^*/+; Ai14/+* and 2) Runx1^CKO^: *bhlhe22*^*Cre*^*/+; Ai14/+; Runx1*^*F/F*^. SGNs were dissociated for single cell transcriptomic profiling as follows: cochleae were extracted from euthanized mice and dissected in chilled Leibovitz’s L-15 buffer containing 30 uM Actinomycin D (ActD). They were then treated with collagenase type IV, followed by papain, both in the presence of 15 µM ActD, for 20 minutes each at 37 °C. Papain was inactivated by passing the cells through ovomucoid as recommended for the Papain Dissociation System (Worthington). Dissociated cells were resuspended in chilled 1x PBS containing BSA and passed through a 35 µm filter to remove cell clumps and undigested tissue. The cell suspension thus obtained was loaded onto a MoFlo Astrios EQ Cell Sorter (Beckman Coulter) equipped with a 100 µm nozzle and 561 nm laser to enrich for tdTomato+ cells. Depending on recovery rate, 3000 to 6000 tdTomato+ cells were loaded along with appropriate Chromium Single Cell 3’ reagents (v3) per well of a Chromium Chip B and processed using a 10x Chromium Controller following manufacturer recommendations. cDNA was amplified via 13 PCR cycles in a BioRad C1000 thermocycler. cDNA libraries containing standard P5 and P7 Illumina paired-end constructs were sequenced in an Illumina NextSeq 500 platform using a 75-cycle high-output kit. Each run of the 10x Chromium Controller consisted of a control and a Runx1^CKO^ sample, each loaded on a separate well. These samples were processed separately until library preparation but pooled for sequencing.

### Statistics

All statistical analysis was done in R. For comparing two independent groups, normality of data distribution within each group was assessed by the Shapiro-Wilk test. If both groups showed normal distribution, Welch’s t-test was performed. Otherwise, the non-parametric Wilcoxon Rank Sum Test or Kolmogorov-Smirnov (KS) Test was applied. Data are presented in the text as mean±SD.

### Bioinformatic analysis

Sequenced reads were demultiplexed to separate by experimental group and aligned to the mouse reference genome (mm10) in a Linux-based high-performance computing cluster at Harvard Medical School. Bioinformatic analysis was performed using custom scripts that utilized Seurat for preliminary processing and various other packages for statistical analysis and graphical representation in the R environment. Principal component analysis (PCA), UMAP embedding, and graph-based clustering were all performed within the Seurat environment.

Supervised classification was performed using SingleCellNet. Ensemble learning-based approaches coerce cells, by design, into one of several possible classes present in training data (i.e., control SGNs), so cases of mixed or unresolved identities absent in the wildtype state are missed. To get around this, a 4^th^ cell class was created by randomly permuting gene expression across all three Type I SGN subtypes, thereby creating a synthetic identity. Any cell that either matched this mixed cell class or exhibited poor resemblance (<50%) to the other pure cell classes were assigned this mixed identity. For supervised classification based on gene subsets, DE genes (among Ia, Ib and Ic SGNs in the Control group) that belonged to the following Gene Ontology groups were used: Functionally relevant (GO:0005216 & GO:0007268), Ion channel (GO:0005216), Chemical synaptic transmission (GO:0007268). List of DE genes of the cell adhesion molecule family were drawn using those identified as CAM genes previously (Földy et al., 2016).

Doublets were detected using the DoubletFinder (McGinnis et al., 2019) package in R which injected artificial doublets into our dataset at a set proportion and subsequently defined each SGN’s neighborhood in gene expression space. Optimal pK value was determined by BC_MVN_ (mean-variance-normalized bimodality coefficient) maximization: the pK parameter was varied over a range before picking one that yielded the highest BC_MVN_ (Fig. S1H). Finally, the proportion of artificial nearest neighbors (pANN) was calculated (Fig. S1I). Cells with the highest pANN values were deemed to be doublets as described previously (Fig. S1J).

### Image acquisition

Tissues probed by *in situ* hybridization and immunohistochemistry assays were imaged using a Leica SP8 point-scanning confocal microscope equipped with HyD and photomultiplier tube (PMT) detectors. RNAscope signals were captured with PMT detectors while immunostained cytoplasmic markers (HuD, tdTomato or YFP) were imaged with HyD or PMT detectors, both using 63x oil-immersion objective at 0.180 micron/pixel *xy* resolution. Whole-mount preparations used for synapse analysis were imaged using PMT detectors and 63x oil-immersion objective (voxel size = 0.901 × 0.901 × 0.299 µm^3^). Frequency maps were generated based on low-magnification views of cochlear pieces using the Measure_Line ImageJ plugin available through the Histology Core at Mass Eye and Ear (Boston, USA). Care was taken not to oversaturate any fluorescent signal intended to be used for intensity-based morphometric quantification. Fluorescent signal meant for generating cell segmentation masks (i.e., hair cell or SGN cytoplasmic markers) were intentionally oversaturated; this was necessary to capture the full volumes of respective cells by compensating for the tendency of the software to draw cell surfaces inside actual cell boundaries.

### Image analysis

#### In situ gene expression assays

Quantification of RNAscope signal in tissue sections was performed by generating counts of fluorescent puncta within SGN soma segmented based on anti-HuD in Imaris (BitPlane). Detection of genetically encoded fluorescent reporter (e.g., tdTomato, YFP) in lineage-tracing experiments was performed similarly, except the Imaris output was binarized to yield ON/OFF status for each SGN. Subsequent data processing and statistical analysis was done in the R environment.

#### Neuronal density in the SG

SGN density was analyzed blind to genotype by following these steps in ImageJ: (1) manually counting neuronal cell bodies in maximum intensity projections of spiral ganglia of cryosectioned cochlea; and (2) measuring ganglion area using the ‘Polygon selections’ tool and the ‘Measure’ function.

#### Synapse count, position and morphometry

A combination of Imaris, MATLAB and R was used to analyze HC-SGN synapses marked by anti-CtBP2 and anti-GluA2: creation of 3D hair cell segmentation masks as well as determination of synaptic element volume and position in image-centric cartesian coordinates (Fig. 4B) was accomplished semi-automatically in Imaris. Membership of synaptic elements to particular HCs was determined automatically in Imaris based on inclusion within a segmented volume. Any Imaris-rendered HC segmentation mask meeting the following criteria were excluded from further analysis: (1) missed portions of the HC and synaptic elements associated with it; and (2) encapsulated >1 HC. References for HC-centric coordinates were then created in ImageJ by manually annotating the following landmarks as singular points in the z-stack for each HC using a custom macro: the back of the cuticular plate (i.e., the side closest to the tallest sterocilia row), the center of the nucleus, mid-point along the bottom of the hair cell. Data generated in Imaris and ImageJ were then processed using custom-written R and MATLAB scripts. Notably, image-centric coordinates (Fig. 4A) corresponding to position of synaptic elements were transformed to HC-centric coordinates (Fig. 4B) with the origin set at the base of each HC and the coordinates rotated to account for the unique tilt of each HC. The latter was achieved by aligning the y-axis of the new coordinate system with a slicer plane (Fig. S3A) that connected the mid-point at the bottom of the HC with the back of its cuticular plate (Fig. S3A, 4B). The slicer plane effectively partitioned each HC into modiolar (facing the modiolus) and pillar (facing the pillar cell) volumes, which enabled assignment of ‘modiolar’ and ‘pillar’ designations to each synaptic element based on their location within the respective volumes. To account for staining intensity variation across tissues, ribbon and glutamate receptor volumes were normalized relative to the respective median volumes in each z-stack as described previously (Liberman et al., 2011). Synaptic elements that did not have a corresponding pre- or post-synaptic element nearby was excluded from our analysis. A small subset of HCs exhibited extremely skewed distribution of modiolar and pillar synaptic elements (e.g., all elements were modiolar); these were excluded from analysis as they may represent immunostaining, slide mounting or segmentation artifacts. Statistical analysis was performed in R. All of these analyses were done blind to genotype.

### Auditory Brainstem Responses (ABR) and Distortion Products Otoacoustic Emissions (DPOAE)

Mice were anesthetized with an intraperitoneal injection of ketamine (100 mg/kg) and xylazine (7.5 mg/kg) and placed on a 37 ºC heating pad for the duration of experiment. Acoustic stimuli were delivered via a custom acoustic system from the Eaton-Peabody Lab (acoustic assembly and custom Labview software available from the Engineering Core of the Eaton-Peabody Laboratories at Massachusetts Eye and Ear, Boston) in a sound-attenuating, electrically shielded room. A dorsoventral incision at the intertragal notch of the pinna exposed the ear canal.

Distortion Products Otoacoustic Emissions (DPOAEs) were recorded for primary tones (frequency ratio f_2_/f_1_ = 1.2, level ratio L_1_ = L_2_ + 10), where f_2_ varied from 5.6 to 32 kHz in half-octave steps and incremented in 10 dB steps from 10 dB to 70 dB. The cubic distortion product 2f_1_-f_2_ was extracted by Fourier analysis of the ear-canal sound pressure after waveform and spectral averaging. DPOAE threshold was defined as the f_1_ level required to produce a DPOAE of 0 dB SPL.

Auditory brainstem responses (ABRs) were recorded via three subdermal needle electrodes: (1) reference electrode placed caudal to the pinna, (2) active electrode in the vertex, and (3) ground electrode near the tail. Stimuli were 5-ms tone-pips with 0.5-ms rise-fall time at frequencies from 5.66-32 kHz delivered in non-alternating polarity at 30 s^-1^. Stimulus levels (SPL) were incremented in 5 dB steps from 30 dB to 80 dB. Cochlear Function Test Suite (CFTS) software (v2.19; Massachusetts Eye and Ear, Boston) amplified (×10^4^), filtered (0.3-3 kHz passband), and averaged with 512 responses at each SPL.

ABR Peak Analysis software (v.1.1.1.9; Massachusetts Eye and Ear, Boston) was used to determine thresholds and measure peak amplitudes and latencies for ABRs. ABR threshold was defined as the SPL at which a reproducible response waveform appeared. Threshold, peaks and troughs were verified by blinded visual inspection. Wave amplitude was defined as the difference between the peak and subsequent trough.

## Supplementary Tables

**Table S1: Differentially expressed genes among Type I SGN subtypes in Control**

Related to Figure 2. Data obtained using the FindMarkers() function in Seurat (R package) after clustering of Type I SGN scRNA-seq profiles. PCT.1 refers to the proportion of cells expressing the gene in the cluster of interest (‘subtype’ column) while PCT.2 refers to the expressing proportion among all other cells in the dataset.

**Table S2: Genes used for various iterations of supervised classification**

Related to Figure 3. Complete list of genes used for each type of supervised classification.

**Table S3: Proportions of SGN subtypes predicted by supervised classification**

Related to Figure 3D. This table contains the proportion of SGN subtypes predicted for each test data set (i.e., all Type I SGNs or those in cluster Ia, Ib or Ic in the UMAP plot in Figure 2D). Data is separated by gene subset used for classification (columns 3-7) as well as genotype (column 2).

**Table S4: Results of statistical analysis of ABR data**

Related to Figure 5. DPOAE threshold, ABR threshold and Peak 1 amplitude were compared at the indicated sound frequency and/or intensity. Actual p values are reported for Welch’s t-test or Kolmogorov-Smirnov (KS) test. *: p<0.05, NS: p>0.05, -: statistical test not performed due to sample size limitation.

## Supplementary Figures

**Figure S1:**
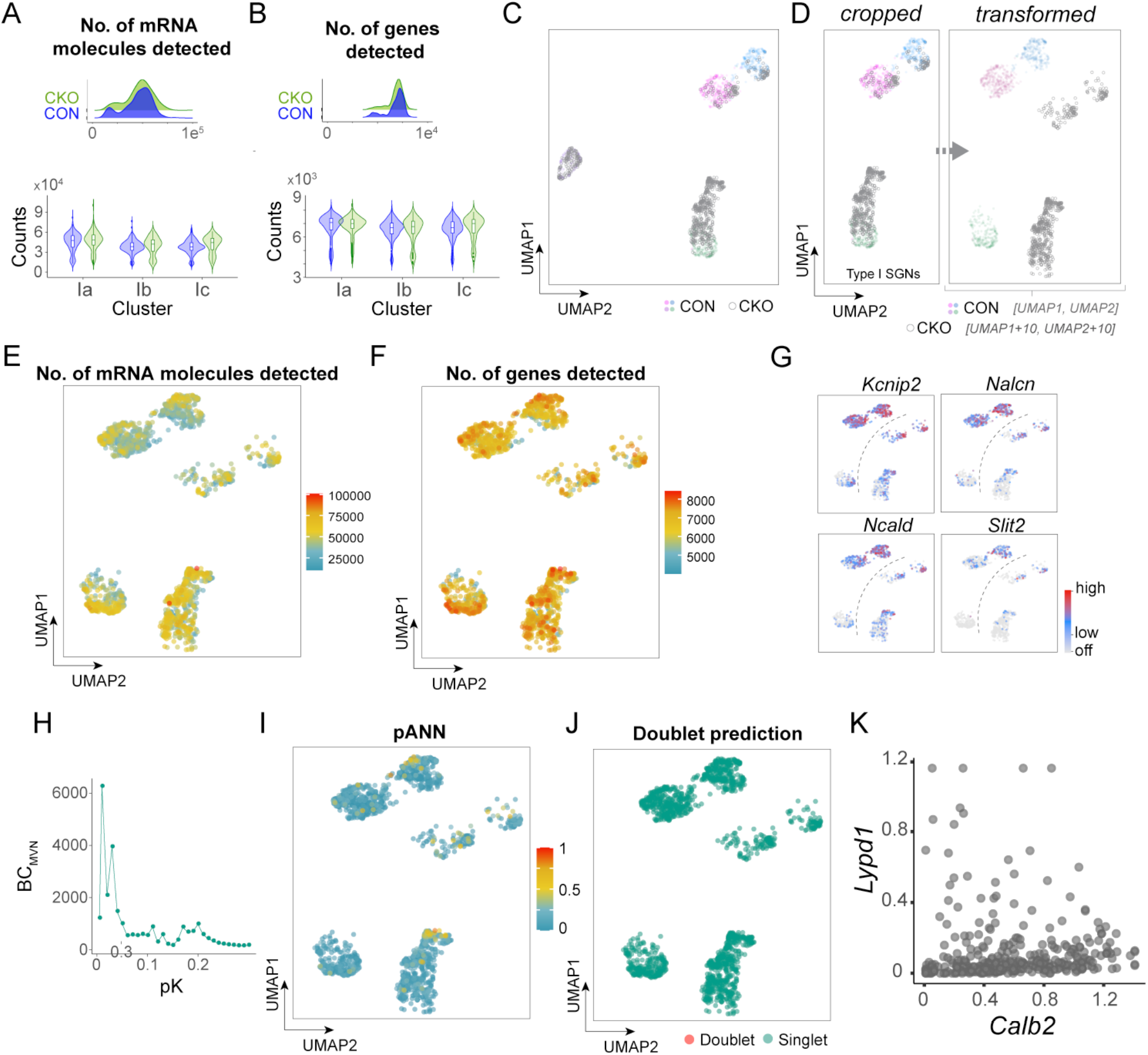
(**A, B**) Group-wise distributions of number (no.) of RNA molecules (A) and no. of genes detected (B) per cell shown as ridge plots (top) and joint violin and box-and-whisker plots split across the three neuronal clusters (bottom). (**C**) Cells from the *Runx1* conditional knock-out (CKO) group showed as gray circles while colored dots represent control (CON) cells in a 2-dimensional UMAP embedding. (**D**) Given our focus on Type I SGNs, the UMAP plot was cropped to show only the portion representing those neurons. To better visualize CKO neurons, their coordinates in the UMAP 2D space were transformed by adding 10 units in both axes. This resulted in spatial separation of otherwise overlapping cells of CON and CKO groups. Such transformed UMAP plots were only used for visualization purposes. (**E, F**) Feature plots of no. of mRNA molecules (E) and no. of genes detected (F) revealed that these metrics did not vary in any systematic way based on cell position in the UMAP space. (G) Feature plot showing expression level of exemplar Ib/Ic or Ic-specific genes that are mostly downregulated in Ia SGNs in CKO. (**H**) Determination of optimal pK value by BC_MVN_ (mean-variance-normalized bimodality coefficient) maximization (see Methods). (**I**) Feature plot showing pANN (proportion of artificial nearest neighbors) values for all cells. (**J**) Results of doublet prediction overlaid on a transformed UMAP plot – no doublets were found. (**K**) Scatterplot of *Calb2* vs. *Lypd1* expression in SGNs, assessed by RNAscope, and showing a view before results of SVM classification were applied.

**Figure S2:**
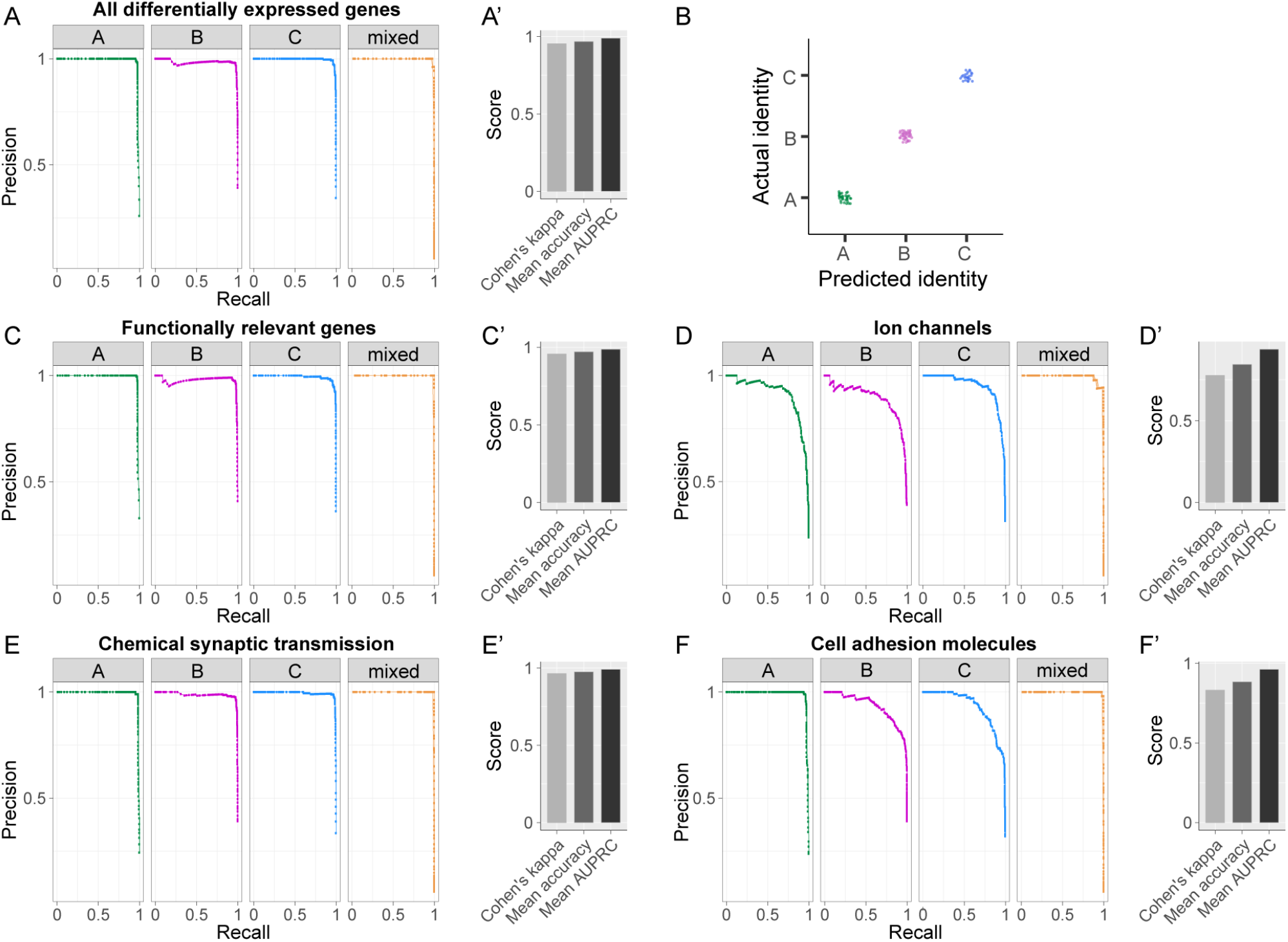
Precision-recall curves for TP-RF classifiers built using various gene subsets (**A**,**C**,**D**,**E**,**F**) indicate capabilities for high precision and high recall, with comparatively poorer performance when only ion channels or cell adhesion molecules were considered. Other metrics of prediction accuracy are shown in (**A’, C’, D’, E’, F’**). Predicted identities were compared against actual identities (determined by unsupervised classification) for supervised classification using all differentially expressed genes (**B**).

**Figure S3:**
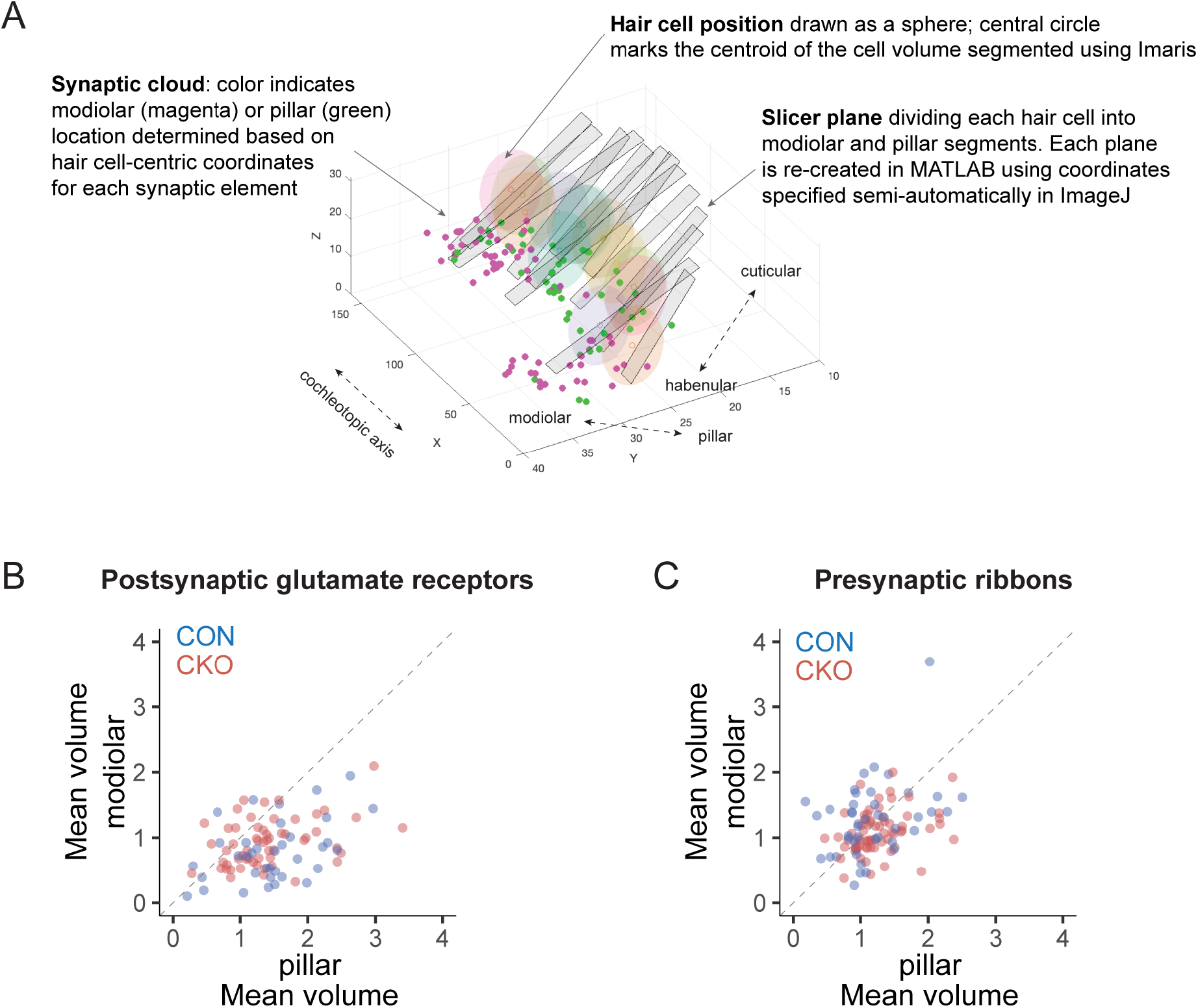
(**A**) Exemplar digital data reconstruction used for comparing HC-SGN synapse properties. Shown is a 3D plot marking HC positions, slicer planes that partition HCs into modiolar and pillar volumes, and synaptic clouds colored to indicate their measured location. (**B, C**) Scatterplots of mean modiolar and pillar volumes of postsynaptic glutamate receptor (B) or presynaptic ribbon (C). Each dot represents an individual HC.

**Figure S4:**
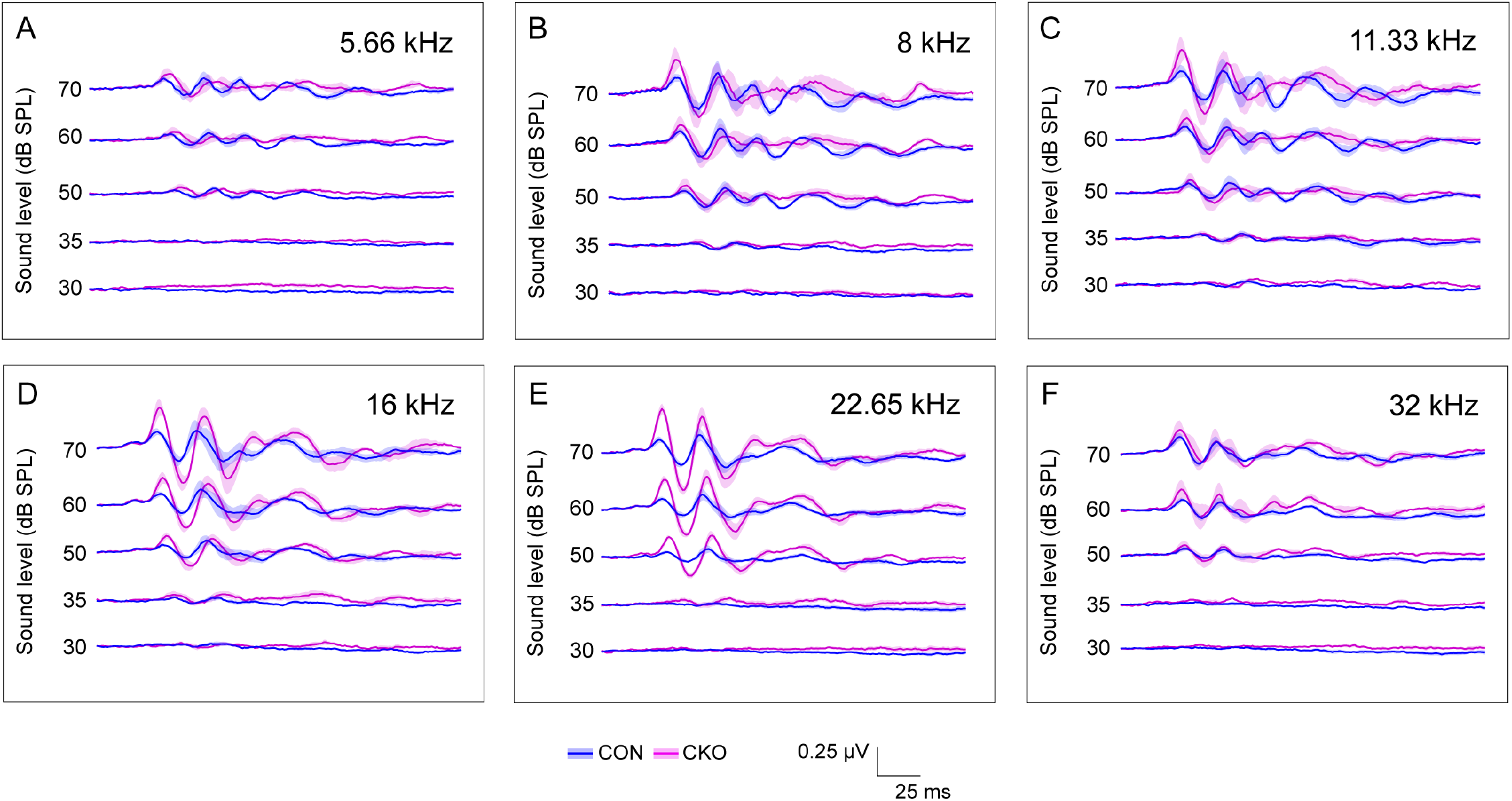
(**A-F**) Averaged (± SEM) auditory brainstem response (ABR) waveforms at select intensities for all tone burst frequencies tested. Comparison across frequencies tested indicate that the strongest and least noisy gains in Peak 1 differences are observed at 16 (**D**) and 22.65 kHz (**E**) regions.

## Notes

### Competing Interest Statement

The authors have declared no competing interest.

